# Early metabolic priming under differing carbon sufficiency conditions influences peach fruit quality development

**DOI:** 10.1101/2020.09.01.278499

**Authors:** Brendon M. Anthony, Jacqueline M. Chaparro, Jessica E. Prenni, Ioannis S. Minas

## Abstract

Crop load management is an important preharvest factor to balance yield, quality, and maturation in peach. However, few studies have addressed how preharvest factors impact metabolism on fruit of equal maturity. An experiment was conducted to understand how carbon competition impacts fruit internal quality and metabolism in ‘Cresthaven’ peach trees by imposing distinct thinning severities. Fruit quality was evaluated at three developmental stages (S2, S3, S4), while controlling for equal maturity using non-destructive near-infrared spectroscopy. Non-targeted metabolite profiling was used to characterize fruit at each developmental stage from trees that were unthinned (carbon starvation) or thinned (carbon sufficiency). Carbon sufficiency resulted in significantly higher fruit dry matter content and soluble solids concentration at harvest when compared to the carbon starved, underscoring the true impact of carbon manipulation on fruit quality. Significant differences in the fruit metabolome between treatments were observed at S2 when phenotypes were similar, while less differences were observed at S4 when the carbon sufficient fruit exhibited a superior phenotype. This suggests a potential metabolic priming effect on fruit quality when carbon is sufficiently supplied during early fruit growth and development. In particular, elevated levels of catechin may suggest a link between secondary/primary metabolism and fruit quality development.

**Highlight:** An investigation of variable carbon supply conditions in peach fruit reveals that early metabolic priming is associated with quality development

## Introduction

Improvement of peach (*Prunus persica* L. Batsch) fruit quality is of critical importance as peach consumption has been in steady decline in recent years due to poor performance with consumers (Minas *et al*., 2018). It is broadly accepted and supported by previous studies that fruit quality can only be improved at the preharvest stage in the orchard, and then maintained during postharvest handling (Crisosto *et al*., 1997; Minas *et al*., 2018). One of the most influential preharvest factors for peach fruit quality is crop load management (e.g. flower/fruitlet thinning) (Minas *et al*., 2018). Advantages of thinning include increased fruit weight and size, higher levels of dry matter content (DMC) and soluble solids concentration (SSC), and overcolor blush coverage (Marini, 2003; Crisosto and Costa, 2008; Minas *et al*., 2018, 2021). Thinning flowers or fruitlets balances the leaf-to-fruit ratio, or the source/sink relationship for photosynthates. This potential yield sacrifice is crucial to reduce competition for photosynthates between fruits, so that the remaining fruit have an adequate pool of carbohydrates at every stage of development enabling them to reach maximum growth and quality at harvest (Grossman and DeJong, 1995).

The main forms of carbon translocated from leaves to fruit are sorbitol and sucrose, via symplastic and apoplastic pathways, to support growth and development in peach (Zhang *et al*., 2004; Morandi *et al*., 2008). Plasma membrane-bound sorbitol transporters (SOTs) take up sorbitol into the cytosol of parenchyma cells where it is rapidly converted to fructose by sorbitol dehydrogenase (SDH) (Zhang *et al*., 2004). Conversely, sucrose, is either directly taken up by sucrose transporters (SUC; SUT) into the cytosol, or degraded to hexoses (e.g. glucose and fructose) by cell wall invertases (CWINVs), which are then transported into the cytosol via hexose transporters (Bologa *et al*., 2003; Zhang *et al*., 2004). Cytosolic hexoses are phosphorylated through the glycolytic pathway and split into triose phosphates, which are finally oxidized to pyruvate (Li *et al*., 2018). Pyruvate can then be transported into the matrix of the mitochondria by a mitochondrial pyruvate carrier to fuel the tricarboxylic acid (TCA) cycle (Li *et al*., 2018; Boeckx *et al*., 2019). Unrestricted availability of these soluble sugars is of critical importance for the maintenance of the fruit’s respiratory metabolic functions, which are vital to producing energy, and assembling carbon-based compounds for use in various biosynthetic processes in primary and secondary metabolism (Boeckx *et al*., 2019).

Similarly to carbon synthesis and utilization, maturation and ripening in climacteric fruit (e.g. peach) are highly regulated processes that initiate with an increase of respiration and release of ethylene (Giovannoni *et al*., 2017). Throughout “on-tree” maturation and ripening of peach, several sensorial and textural changes occur, such as increasing DMC and SSC, pigment accumulation, flesh softening, and aromatic volatilization; all parameters that improve organoleptic characteristics and consumer satisfaction (Crisosto and Costa, 2008; Minas *et al*., 2018). Preharvest factors that affect carbohydrate starvation/competition in the growing fruit (e.g. girdling, leaf-to-fruit ratio, crop load) can severely alter the regulation of the primary and secondary metabolism, influencing the maturation/ripening processes and internal fruit quality (Nardozza *et al*., 2019). Importantly, preharvest factor comparisons performed on fruit of unequal maturity are potentially confounded by the maturation status of the fruit. Therefore, to truly evaluate the impact of preharvest conditions on fruit physiology and metabolism it is critical that comparisons be made on fruit of equal maturity (Minas *et al*., 2018).

Understanding how preharvest factors impact fruit internal quality (DMC, SSC) and maturity (flesh firmness, FF) is essential, but can be very time-consuming and expensive with traditional destructive techniques (Minas *et al*., 2018). Furthermore, given the variability of fruit on a tree, or within an orchard, these narrow approaches fail to capture the totality of quality development in the field (Carlomagno *et al*., 2004). Near infrared spectroscopy (NIRS) is an alternative non-destructive technique that has been successfully utilized to accurately assess different quality attributes in peach (Minas *et al*., 2021). In addition, in recent years, a new index (index of absorbance difference, I_AD_) has been developed to characterize physiological changes (e.g. chlorophyll degradation) that occur during peach maturation and ripening (Ziosi *et al*., 2008; Costa *et al*., 2009). This index effectively describes the physiological maturity of peach and has been integrated into open-source handheld Vis-NIRS sensors to enable simultaneous and rapid evaluation of peach fruit internal quality and physiological maturity (Minas *et al*., 2021). This technology can be paired with molecular tools to better understand regulation differences in fruit, as it can enable rapid maturity control assessments.

Given the complexity of the biological processes associated with fruit maturation, ripening, and quality development, the use of large-scale molecular tools (“omics”) that allow for the global evaluation of metabolic pathways within the fruit is particularly important (Molassiotis *et al*., 2013). Non-targeted metabolomics is an approach that can be utilized in horticultural research to advance our understanding of the impact of preharvest factors on regulatory networks and biochemical interactions in plants (Guy *et al*., 2008). Fruit is the most metabolite-rich plant organ, and thus metabolomics offers an opportunity to investigate the metabolic processes involved in quality development (Monti *et al*., 2016). Thus, regulation of the central carbon metabolism through manipulation of preharvest factors is not only essential for fleshy fruit carbohydrate accumulation (e.g. sugars and organic acids), growth, development, maturation, and ripening, but it is also key to fruit taste, flavor and quality (Cirili *et al*., 2016; Li *et al*., 2018; Nardozza *et al*., 2019). In addition, the accumulation of specific sugars, organic acids or secondary metabolites at different phases of fruit development as a response to stress or preharvest conditions might be related to signaling or priming procedures (Tohge *et al*., 2014).

Previous studies that have been conducted to better understand metabolic regulation in peach have primarily evaluated postharvest factors and cold storage disorders (Lauxmann *et al*., 2014; Bustamante *et al*., 2016; Tanou *et al*., 2017; Santin *et al*., 2018; 2019; Monti *et al*., 2019; Lillo-Carmona *et al*., 2020), along with maturation/ripening physiology, environmental conditions and pest tolerance (Lombardo et al., 2011; Capitani et al., 2012; Monti *et al*., 2016; Karagiannis *et al*., 2016). Characterizing fruit metabolic shifts in response to preharvest factors is key to fruit quality improvement, as several of these compounds contribute to organoleptic characteristics that correlate with consumer preference. Many studies have reported effects of various preharvest and orchard factors on tree fruit quality (Minas *et al*., 2018, Musacchi and Serra, 2018). However, despite the enormous advances in “omics” sciences, few experiments have addressed the influence of these preharvest factors on fruit quality at the metabolic level (Monti *et al*., 2016; Serra *et al*., 2018; Michailidis *et al*., 2020). Furthermore, it is important to highlight that these previous studies did not control for equal maturity when analyzing fruit coming from various preharvest conditions. Therefore, to truly understand how these factors influence both fruit quality and metabolism, comprehensive experimental approaches that control for maturity amongst comparisons are critical pre-requisites (Minas *et al*., 2021).

The goal of this study was to evaluate the true impact of crop load management, a major preharvest factor, on peach quality using phenotypic and metabolomic assessments of fruit of equal maturity. Our experimental approach enabled a detailed physiological characterization of photosynthate availability reflective of variable fruit-to-fruit competition conditions (i.e. thinning severities) during peach fruit growth and development. Using non-destructive NIRS technology (to control for fruit maturity) and non-targeted metabolite profiling to evaluate the impact of carbon supply, we identified metabolic shifts associated with fruit quality priming/suppression under carbon sufficiency/starvation.

## Materials and Methods

### Plant material and experimental approach

Fifteen peach (*Prunus persica* L. Batsch.) trees of a late-ripening cultivar (‘Cresthaven’) and of uniform size and health were selected from Colorado State University’s experimental orchard at the Western Colorado Research Center-Orchard Mesa, Grand Junction, CO (39°02’31.3”N, 108°27’56.8”W). The trees were nine years old, grafted on ‘Lovell’ rootstock, and trained to an open vase system at a planting density of 509 trees ha^-1^ (4 × 5 m spacing). Fifty-two days after full bloom (DAFB: April 9, 2016) five thinning treatments were administered to shift the photosynthate source:sink ratio and create distinct carbon supply levels. Each treatment was performed on three replicate trees. The treatments were designed to achieve varying levels of photosynthate sufficiency for the growing fruits and included leaving trees unthinned (control) or by thinning and spacing the remaining fruit every 5, 10, 15 and 30 cm from each other on the fruiting shoots (Fig. 1A). Trees were thinned after “June drop” during early stage 2 (S2) of peach fruit development to ensure no further natural fruit abscission would occur and to maintain the spacing and predetermined levels of carbon competition amongst the remaining fruit. The trees were managed according to industry standards and practices.

**Fig. 1.**
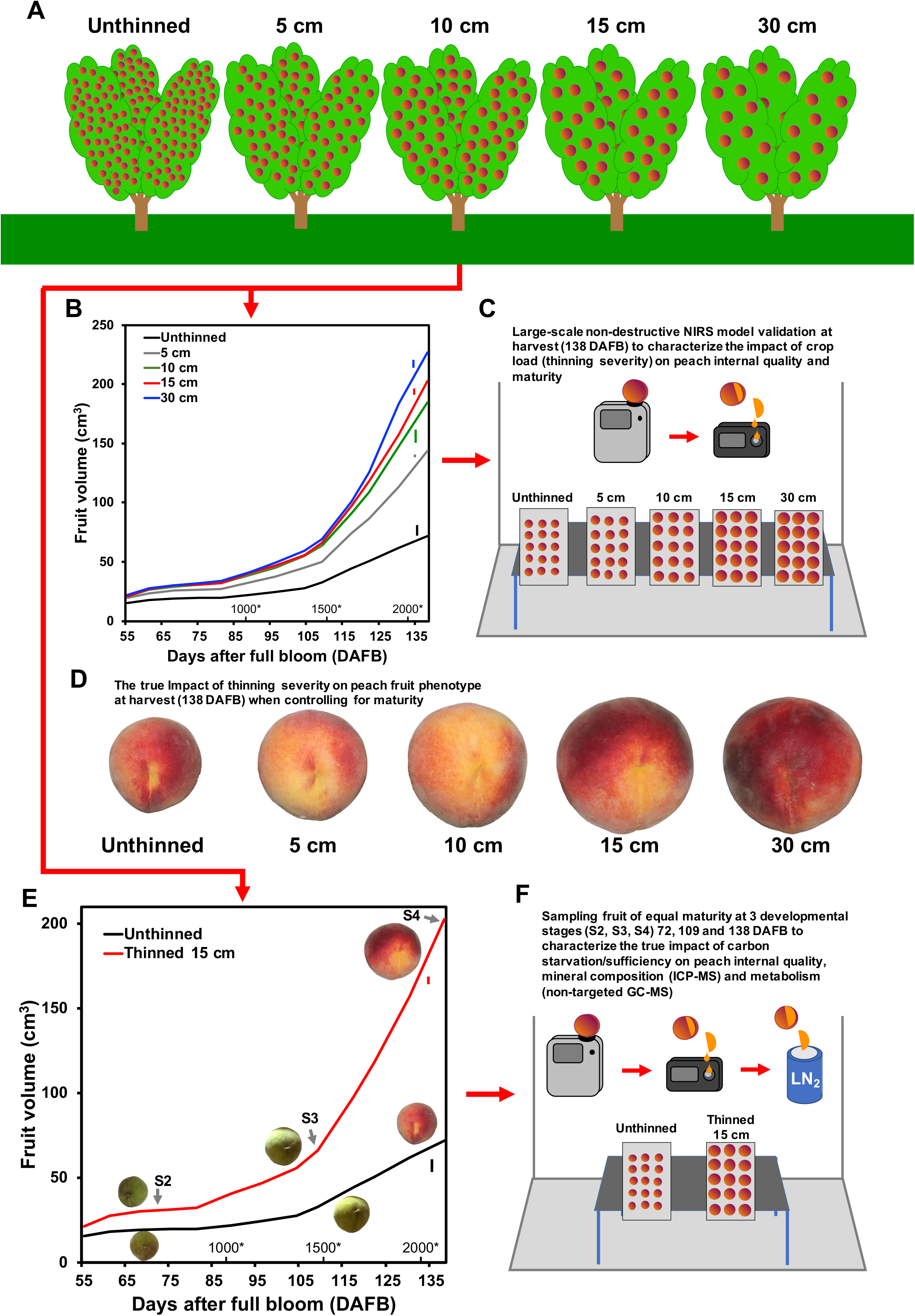
Carbon supply manipulation to assess peach fruit quality, elemental composition and metabolism. A carbon supply manipulation study was conducted to understand the true impact of thinning severity on fruit quality, along with how fruit quality development is linked with ionome and metabolome. Trees were thinned to different fruit spacing distances (5, 10, 15 and 30 cm) and compared with unthinned control trees (A). Thinning was conducted 52 days after full bloom (DAFB). Fruit from these trees were evaluated non-destructively throughout the growing season for volumetric growth. Fruit growth assessment time is indicated in DAFB and growing degree days (GDD) with asterisks above. Vertical colored bars represent the least significant difference (LSD) per treatment based on Tukey’s HSD test (*P*<0.05) (B). Fruit coming from distinct thinning severities were assessed for quality and physiological maturity at harvest (S4; 138 DAFB) with a non-destructive near infrared spectrometer (NIRS), along with destructive analyses for model validation (C). Fruit were evaluated for their phenotype while controlling for equal maturity to understand the true impact of thinning severity on quality across all treatments (D). Two thinning treatments were selected for further biological analyses: the carbon starved, unthinned treatment, and the optimal thinned at 15 cm fruit spacing treatment, carbon sufficient. Volumetric growth is displayed between the unthinned and thinned treatments across growth and development (S2-S4), while controlling for equal maturity. Vertical bars represent LSD according to Tukey’s HSD test (*P*<0.05) (E). At each developmental stage (S2, S3, S4) 72, 109 and 138 DAFB, respectively, fruit were harvested to characterize the true impact of carbon supply on peach internal quality, which allowed for further investigation into the fruit ionome and metabolome. Samples were immediately frozen with liquid nitrogen (LN_2_) and lyophilized at each developmental stage to quench the metabolism, until molecular analyses (ICP-MS and non-targeted GC-MS) could be conducted (F).

Trunk cross sectional area (TCSA) was measured in the spring to enable the selection of uniform trees, and again in the fall (postharvest) to assess tree growth responses to treatments throughout the growing season and to calculate crop load (fruit · cm^-2^ TCSA) (Table 1). Fruit growth (diameter, mm) was monitored in 10 fruit per experimental tree the day prior to thinning, immediately after thinning and every week the rest of the growing season until harvest, using a fruit lasso. Peach volume (cm^3^) was calculated as was previously described (Minas *et al*., 2015; Fig. 1B, E).

**Table 1.** Effect of thinning severity on peach tree physiology and fruit external quality attributes. The impact of thinning severity on trunk cross sectional area (TCSA) (cm^2^), number of fruit per tree, crop load (fruit per cm^2^ of TCSA), yield (kg tree^-1^), average (avg.) fruit weight (g), fruit diameter (mm), overcolor (%), number of flowers/branch, and fruit set (%) in ‘Cresthaven’ peach. ‘Cresthaven’ peach trees were thinned to various levels (5, 10, 15 and 30 cm) and compared with unthinned control trees. Mean values ± S.E. are displayed. Means followed by the same superscript letter are not statistically different according to Tukey’s HSD test (*P*<0.05).

| Thinning treatment | TCSA<sup>†</sup> (cm<sup>2</sup>) | No. of fruit per tree | Crop load (no. of fruit/cm<sup>2</sup> of TCSA) | Yield (kg/tree) | Production (MT/ha) | Avg. fruit weight (g) | Diameter (mm) | Overcolor (%) | Return bloom (no. of flowers per branch<sup>#</sup>) | Return fruit set<sup>‡</sup> (%) |
| --- | --- | --- | --- | --- | --- | --- | --- | --- | --- | --- |
| Unthinned | 62.0 ± 4.1 | 731 ± 51<sup>a</sup> | 11.8 ± 0.27<sup>a</sup> | 61.5 ± 6.9<sup>a</sup> | 31.2 ± 3.5<sup>a</sup> | 84.6 ± 9.3<sup>c</sup> | 57 ± 0.4<sup>e</sup> | 64 ± 1.0<sup>b</sup> | 9 ± 1<sup>b</sup> | 10 ± 5<sup>c</sup> |
| 5 cm | 60.9 ± 0.3 | 346 ± 43<sup>b</sup> | 5.7 ± 0.72<sup>b</sup> | 53.6 ± 3.6<sup>a</sup> | 27.2 ± 1.8<sup>a</sup> | 157.4 ± 10.4<sup>b</sup> | 69 ± 0.3<sup>d</sup> | 59 ± 1.0<sup>c</sup> | 18 ± 1<sup>a</sup> | 23 ± 5<sup>bc</sup> |
| 10 cm | 61.0 ± 5.1 | 220 ± 45<sup>bc</sup> | 3.6 ± 0.62<sup>c</sup> | 40.5 ± 9.7<sup>ab</sup> | 20.6 ± 4.9<sup>ab</sup> | 181.5 ± 6.5<sup>b</sup> | 75 ± 0.3<sup>c</sup> | 67 ± 0.8<sup>b</sup> | 20 ± 1<sup>a</sup> | 30 ± 5<sup>b</sup> |
| 15 cm | 63.8 ± 3.3 | 169 ± 21<sup>c</sup> | 2.6 ± 0.27<sup>cd</sup> | 38.5 ± 4.0<sup>ab</sup> | 19.5 ± 2.0<sup>ab</sup> | 228.6 ± 7.6<sup>a</sup> | 78 ± 0.3<sup>b</sup> | 68 ± 0.9<sup>ab</sup> | 20 ± 1<sup>a</sup> | 56 ± 5<sup>a</sup> |
| 30 cm | 71.9 ± 10.0 | 111 ± 5<sup>c</sup> | 1.6 ± 0.18<sup>cd</sup> | 27.8 ± 1.6<sup>b</sup> | 14.1 ± 0.8<sup>b</sup> | 249.9 ± 13.7<sup>a</sup> | 80 ± 0.4<sup>a</sup> | 71 ± 0.8<sup>a</sup> | 21 ± 1<sup>a</sup> | 34 ± 6<sup>b</sup> |
| Significance | ns | *** | *** | ** |  | *** | *** | *** | *** | *** |

ns, \*, \*\*, \*\*\* indicate no significance or significant at p-values of <0.05, 0.01, or 0.0001
<sup>†</sup>trunk cross sectional area
<sup>#</sup>n=3 branches; quantified at full bloom on 22 March 2017
<sup>‡</sup>fruit set calculated as: number of fruit/number of flowers x 100 on 9 May 2017

Yield data were collected at the conclusion of the experiment (August 25) when fruit were harvested at the pre-climacteric commercial harvest maturity (stage 4I, referred to as S4). The fruit was counted and then weighed to determine yield (kg · tree^-1^), while the average fruit fresh weight (FW, g) was calculated. Crop load was calculated and expressed as fruit no. · cm^-2^ of TCSA (Table 1). Fifteen fruit from each experimental tree (n=45 per treatment) were assessed destructively and non-destructively to evaluate the impact of crop load on peach maturity and internal quality at harvest (Table 2; Fig. 1C).

**Table 2.**
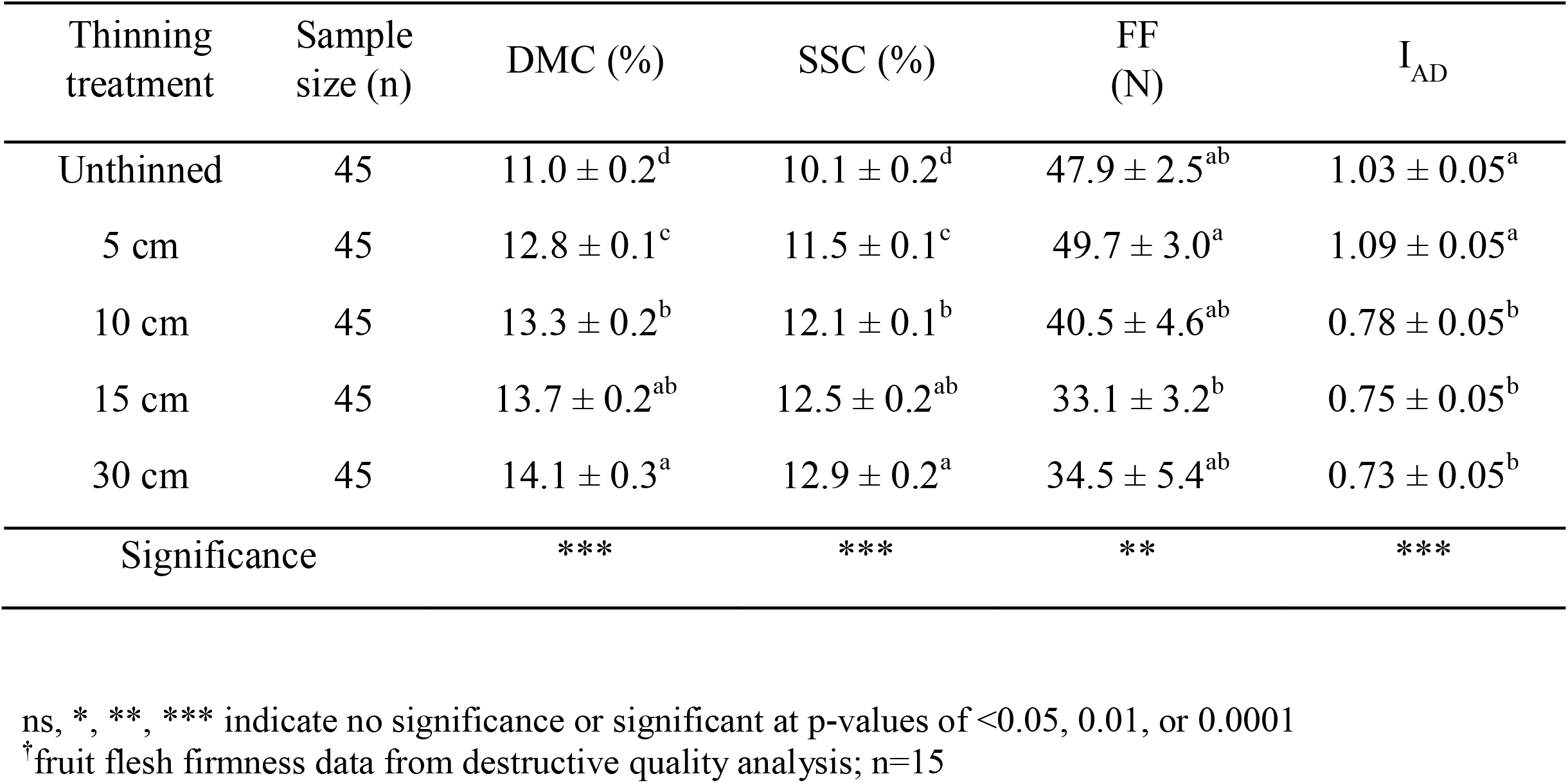
Effect of thinning severity peach fruit internal quality and maturity. The impact of thinning severity on fruit internal quality and maturity, assessed with a non-destructive NIRS sensor across all treatments, in ‘Cresthaven’ peach. ‘Cresthaven’ peach trees were thinned to various levels (5, 10, 15 and 30 cm) and compared with unthinned control trees. Peach fruits (n=45) were assessed non-destructively for internal quality and maturity by measuring for dry matter content (DMC) and soluble solids concentration (SSC) along with flesh firmness (FF) and index of absorbance difference (I_AD_). Due to a lack of accuracy in non-destructive estimation of FF, destructive values (n=15) are displayed. Performance of the non-destructive prediction models (DMC, SSC and I_AD_) is presented in Fig. 2A, B, and C. Mean values ± S.E. are displayed. Means followed by the same letter are not statistically different according to Tukey’s HSD test (*P*<0.05).

| Thinning treatment | Sample size (n) | DMC (%) | SSC (%) | FF (N) | I<sub>AD</sub> |
| --- | --- | --- | --- | --- | --- |
| Unthinned | 45 | 11.0 ± 0.2<sup>d</sup> | 10.1 ± 0.2<sup>d</sup> | 47.9 ± 2.5<sup>ab</sup> | 1.03 ± 0.05<sup>a</sup> |
| 5 cm | 45 | 12.8 ± 0.1<sup>c</sup> | 11.5 ± 0.1<sup>c</sup> | 49.7 ± 3.0<sup>a</sup> | 1.09 ± 0.05<sup>a</sup> |
| 10 cm | 45 | 13.3 ± 0.2<sup>b</sup> | 12.1 ± 0.1<sup>b</sup> | 40.5 ± 4.6<sup>ab</sup> | 0.78 ± 0.05<sup>b</sup> |
| 15 cm | 45 | 13.7 ± 0.2<sup>ab</sup> | 12.5 ± 0.2<sup>ab</sup> | 33.1 ± 3.2<sup>b</sup> | 0.75 ± 0.05<sup>b</sup> |
| 30 cm | 45 | 14.1 ± 0.3<sup>a</sup> | 12.9 ± 0.2<sup>a</sup> | 34.5 ± 5.4<sup>ab</sup> | 0.73 ± 0.05<sup>b</sup> |
| Significance |  | *** | *** | ** | *** |

ns, \*, \*\*, \*\*\* indicate no significance or significant at p-values of <0.05, 0.01, or 0.0001
<sup>†</sup>fruit flesh firmness data from destructive quality analysis; n=15

To determine potential implications of carbon competition on tree mineral status 100 leaves were sampled mid-season (July 20) from the middle portion of several one-year-old shoots from each experimental tree (Gavlak *et al*., 2003). These leaves were then washed, dried, and shipped to Ward Laboratories (Kearney, NE, USA) for elemental analyses. Macro- and micro-nutrients assessed included: N %, P %, K %, S %, Ca %, Mg %, Zn ppm, Fe ppm, Mn ppm, Cu ppm, B ppm and Mo ppm (Supplementary Table S1 available at *JXB* online). To assess the impact of carbon supply on return bloom next season, five shoots per experimental tree, the number of flowers, and their length were measured at full bloom in spring (March 22, 2017). The fertility index was then calculated by dividing the number of flower buds by the length of the shoot (cm). Later in spring (May 9), the fruit were counted, and fruit set (%) was calculated (Table 1).

To characterize the true impact of carbon supply on peach quality development without the confounding impact of maturation status, two crop load treatments were selected to be studied in detail during peach growth and development on fruit of equal maturity (Fig. 1E, F). Specifically, the unthinned treatment which represents a carbon “starved” condition due to the high demand and competition between fruits, and the 15 cm fruit-to-fruit spacing (thinned) treatment, which represents an adequate and sufficient carbon supply throughout development due to reduced photosynthate competition were chosen. Five fruit of equal maturity (determined non-destructively using NIRS) per experimental tree (n=15 per treatment) were evaluated for quality (non-destructively and destructively as described below) at three development stages: stage 2 (S2, 72 DAFB), stage 3 (S3, 109 DAFB) and stage 4 (S4, 138 DAFB) (Fig. 1E, F). Three biological replicates consisting of five homogenized fruit mesocarp samples of equal maturity from these two treatments at the three developmental stages were sampled, flash frozen (quenched) with liquid nitrogen and stored at -80 °C (Fig. 1F) until analysis by inductively coupled plasma mass spectrometry (ICP-MS) and gas chromatography mass spectrometry (GC-MS). Prior to analysis, peach mesocarp samples were freeze dried with the lyophilizer (Freezone 4.5, Labconco, Kansas City, MO, USA) at -40 °C for 12 h. Finally, samples were pulverized into a powder with a bead beater (Bullet Blender Storm, Next Advance, Troy, NY, USA) for five minutes. These samples were then kept in -20 °C until digestions and/or extractions could take place.

### Fruit quality analyses

Fruit were assessed for quality, non-destructively, using an “open-type” near-infrared handheld spectrometer (F-750 Produce Quality Meter, Felix Instruments Inc., Camas, WA, USA). The F-750 produce meter was calibrated to non-destructively estimate ‘Cresthaven’ fruit quality based on dry matter content (DMC, %) and soluble solids concentration (SSC, %) (Minas *et al*., unpublished) and physiological maturity based on the index of absorbance difference (I_AD_) with a single scan as previously described (Minas *et al*., 2021). This approach allows for rapid assessment of fruit quality and maturity “on-tree,” to assure that comparisons among treatments and sampling for large-scale analytical methodologies are based on fruit of equal maturity. The non-destructive estimations were validated at each stage with actual destructive data, by scanning and measuring for each parameter from the same side of the fruit as described for field validations (Minas *et al*., 2021; Fig 1).

Physiological maturity measurements were validated using a factory calibrated Vis-NIR spectrometer (DA meter, Sinteleia SRL, Bologna, Italy), which uses the non-destructive metric, I_AD_. The DA meter assesses the absorbance difference (I_AD_=A670 nm-A720 nm) at the specific wavelengths that chlorophyll is absorbing light beneath the fruit surface, which provides an estimate of fruit physiological maturity (i.e. background color) (Ziosi *et al*., 2008; Costa *et al*., 2009; Spadoni *et al*., 2016). Fruit were scanned with the F-750 and DA meter on the sun exposed and shaded side of the fruit, along the equatorial region, and these values were averaged.

Dry matter content measurements were validated destructively by cutting out 25-mm diameter peach mesocarp cylinders with the skin removed using a cork borer. Peach mesocarp cylinders were weighed for fresh weight (FW) using a digital scale (TC-204, Denver instruments, Arvada, CO, USA) and then moved into a forced-air oven (VWR Oven F Air 104 L, VWR, Radnor, PA) at 65 °C to dehydrate, until samples reached a stable weight (∼three days). Dry weight (DW) of the dehydrated peach samples were then measured. Dry matter content was then determined by the mass difference between the FW and DW of each peach sample, as previously described (Minas *et al*., 2021). For SSC (%) measurement validations, similar peach mesocarp cylinders (that were cut from the opposite sides of DMC samples) were juiced through a garlic press into a digital refractometer (PR-32α, Atago, Tokyo, Japan) as previously described (Minas *et al*., 2021).

Additional quality analyses were conducted at each developmental stage, which included parameters such as: skin color (hue angle, *h°*), overcolor blush (%), fruit size (mm), individual fruit fresh weight (FW, g), flesh firmness (FF, N) and titratable acidity (malic acid, %) as previously described (Minas *et al*., 2021).

### Ionomic analysis using inductively coupled plasma mass spectrometry (ICP-MS)

The lyophilized, pulverized, and homogenized samples were randomized into standard 75 mL microwave vessels (PerkinElmer, Waltham, MA, USA). Each vessel received 125 mg of peach mesocarp tissue placed inside of a teflon weighing cup. After the peach tissue was placed into the vessel, 8 mL of doubly distilled nitric acid [HNO_3_, 70% by volume solution spiked with an Indium (In) internal standard] was added and the vessels were left to react for 15 min. An additional 2 mL of ultra-trace grade hydrogen peroxide (H_2_O_2_, 30% by volume solution) was added to each vessel and left to react for an additional 15 min. The seals and rupture discs were then placed onto the vessels, the caps were screwed on and they were loaded into the microwave (Titan MPS Microwave Sample Preparation System, PerkinElmer, Waltham, MA, USA). Samples were processed in random batches of 15 plus a blank (capacity of the Titan MPS system). The preset method (provided by the manufacturer) for “dried fruit” was utilized (Supplementary Table S2). After the completion of the digestion procedure, the vessels were removed and placed into a refrigerator (4 °C) for 30 min to cool. Once cooled, the contents from each vessel was poured into a 50 mL conical falcon tube (Thermo Fisher Scientific, Waltham, MA, USA) and the volume was raised to 15 mL total with 18.2 MΩ ultrapure water (Milli Q Direct, Millipore Sigma, Bedford, MA, USA). This first dilution was then followed by a second dilution, in which 1 mL of the diluted and digested sample from each 50 mL conical falcon tube was then pipetted into a 15 mL conical falcon tube (Thermo Fisher Scientific, Waltham, MA, USA) and again raised to a total volume of 15 mL with Milli-Q water. The final solution contained an internal standard of 20 nL · L^-1^ of In and 2.5% nitric acid.

ICP-MS was performed using the NexION 350D (PerkinElmer, Waltham, MA, USA). In each sample, elemental concentrations of Arsenic (As), Aluminum (Al), Barium (Ba), Boron (B), Cadmium (Cd), Calcium (Ca), Chromium (Cr), Cobalt (Co), Copper (Cu), Iron (Fe), Lead (Pb), Lithium (Li), Magnesium (Mg), Manganese (Mn), Molybdenum (Mo), Nickel (Ni), Phosphorous (P), Potassium (K), Selenium (Se), Sodium (Na), Strontium (Sr), Sulfur (S), Vanadium (V), Tungsten (W), and Zinc (Zn) were measured. Digested samples were injected using a prepFAST SC-2 (Elemental Scientific, Omaha, Nebraska) autosampler. Samples were nebulized using a PFA-ST (Elemental Scientific) nebulizer into a quartz cyclonic spray chamber that was kept at 4 °C via a PC3 peltier cooler (Elemental Scientific). Li, Be, B, Na, P, S, Mg, K, Ca, W, As, and Pb were measured in standard mode. Cd, Se, and As were measured in dynamic reaction (DRC) mode using oxygen as the reactive gas. Al, V, Cr, Mn, Fe, Co, Ni, Cu, Zn, Sr, Mo, and Ba were measured in DRC mode using ammonia as the reactive gas. Prior to analysis, the ICP-MS underwent a daily tune, ensuring that the torch was aligned for maximum In signal. The nebulizer gas flow was also optimized for maximum In signal intensity while also ensuring that oxide formation (CeO^+^:Ce^+^) was kept below 0.02. Additionally, the quadrupole ion deflector (QID) was optimized for maximum signal across the mass range by monitoring Li, Mg, In, Ce, Pb, and U. Furthermore, the formation of doubly charged species was limited by ensuring that Ce^++^:Ce^+^ ratio was below 0.03. A calibration curve was created by evaluating 7 dilutions of a multi-element stock solution, created by a mixture of single element stock standards (Inorganic Ventures, Christiansburg, VA, USA) (Chaparro *et al*., 2018). Calibration curve was matrix matched to the samples, 2.5% HNO_3_ and 20 ppb In. For instrument drift correction, internal standard solutions consisting of ^6^Li, Rh and Ir was added to each sample via the autosampler. Quality control samples were generated by pipetting 1 mL from each sample into a pooled QC and analyzed after every 6 samples.

Ionomic data processing was conducted as previously (Chaparro *et al*., 2018). Data was processed using Excel (Microsoft, Redmond, WA, USA). Each element analyzed was corrected with an internal standard and subsequently underwent drift correction and minimizing the coefficient of variation (CV) of the QC samples identified how each element should be corrected. Samples were then corrected for the appropriate dilution factor. Limits of detection (LOD) and limits of quantification (LOQ) were calculated as 3 times or 10 times the standard deviation of the blank divided by the slope of the calibration curve respectively. Final element concentrations are reported as µg/g of freeze-dried peach mesocarp.

### Non-targeted metabolite profiling using gas chromatography mass spectrometry (GC-MS)

Metabolite extraction was conducted by weighing out 25 mg of each freeze-dried peach mesocarp sample and placing them into clean 2 mL autosampler glass vials (VWR, Radnor, PA, USA). One mL of 80% by volume LC-MS grade methanol (MeOH) in water solution was then added to each vial and vortexed on a plate shaker (Fisherbrand™ Analog Multitube Vortexer, ThermoFisher Scientific, Waltham, MA, USA) at 4 °C at max speed (2500 rpm) for 2 hours (h). Samples were then held at -80 °C for 1 h followed by centrifugation for 25 min at 3500 rpm at 4 °C. The supernatant was extracted (∼800 µL) without disturbing the pellet and pipetted into new a 2 mL autosampler vial. A pooled quality control (QC) solution was created by transferring 10 µL of each sample into a separate glass vial. Fifteen µL of each sample was transferred to another set of glass vials, centrifuged for 2 min at 3500 rpm and then dried under N_2_ (g) for 30 min. Dried samples were stored at -80 °C until derivatization. Derivatization (methoximation and silylation) took place immediately prior to running the samples. Dried down samples were allowed to warm to room temperature and then re-suspended in 50 µL of methoxyamine HCl (prewarmed to 60 °C) and centrifuged for 30 sec. Samples were then incubated at 60 °C for 45 min, followed by a brief vortex, sonication for 10 min and an additional incubation at 60 °C for 45 min. Following this, the samples were centrifuged before receiving 50 µL of N-Methyl-M (trimethylsilyl) trifluoroacetamide (MSTFA) + 1% trimethylchlorosilane (TMCS) (ThermoFisher Scientific, Waltham, MA, USA), briefly vortexed and incubated at 60 °C for 40 min, as described previously (Chaparro *et al*., 2018). Samples were loaded (∼ 80 µL) into glass inserts within glass autosampler vials and centrifuged for 30 sec prior to GC-MS analysis.

GC-MS analysis was performed using the Clarus 690 GC coupled to a Clarus SQ 8S Mass Spectrometer (PerkinElmer, Waltham, MA, USA). Metabolites were separated with a 30 m TG-5MS column (Thermo Scientific, 0.25 mm i.d. 0.25 μm film thickness). The GC program began at 80 °C for 0.5 min and ramped to 330 °C at a rate of 15 °C per minute and ended with an 8 min hold at a 1 mL · min^-1^ helium gas flow rate. The inlet temperature was held at 285 °C and the transfer line was held at 260 °C. Masses between 50-620 m/z were scanned at four scans/sec after electron impact ionization. QC injections were analyzed after every 6^th^ sample and were used to control for and detect analytical variation.

Metabolomic data processing was conducted as previously described (Chaparro *et al*., 2018). GC-MS files were converted to .cdf format and processed by XCMS in R (Smith *et al*., 2006; R Core Team, 2015; Mahieu *et al*., 2016). All samples were normalized to the total ion current (TIC). RAMClustR was used to deconvolute peaks into spectral clusters for metabolite annotation (Broeckling *et al*., 2014). RAMSearch (Broeckling *et al*., 2016) was used to match metabolites using retention time, retention index and matching mass spectra data with external databases including Golm Metabolome Database (Hummel *et al*., 2007; Hummel *et al*., 2013) and NIST (Broeckling *et al*., 2016).

### Statistical analyses

All parameters were assessed for statistical differences between thinning treatments using proc-GLM in SAS (SAS Inc., Cary, NC, USA). The effect of thinning and developmental stage on physiological characteristics, elemental concentrations and mean peak area of metabolites were assessed for significance with either an one-way ANOVA (thinning or developmental stage) or a two-way ANOVA (thinning x developmental stage) (*P*<0.05). Tukey mean comparisons were used to assign different lettering groups where the model was significant at *P*<0.05. Principal component analyses (PCA) was performed on physiological, ionomic, and metabolomic data using SIMCA (Umetrics, Umea, Sweden). Ions, metabolites and fruit physiological characteristics were assessed in R to conduct correlations (using spearman’s rank) and hierarchical clustering using cor, aov, and hclust functions and to develop heat maps using the corrplot package (Wei *et al*., 2010; R Core Team, 2015; Turner *et al*., 2016). Heat maps, PCAs, and graphs were visualized using Prism v8.2.1 (Graph Pad Inc., San Diego, CA, USA).

## Results

### Crop load affected fruit size, quality and maturity at harvest

The various thinning treatments provided a significant impact on the number of fruit per tree, crop load (fruit · cm^-2^ of TCSA) and yield (kg · tree^-1^), although these differences were minimized between the 10 and 30 cm fruit spacing treatments. Final crop loads on trees of similar size, as expressed by TCSA, ranged from 1.6 to 11.8 fruits · cm^-2^ of TCSA (Table 1). Average fruit diameter increased linearly with increasing thinning severity, with a 40% increase, from the unthinned control to extreme thinning (30 cm fruit spacing) (Table 1; Supplementary Fig. S1). A large percentage of fruit (96%) in the extreme crop load (unthinned) were smaller than 70 mm in diameter, whereas the 15 and 30 cm treatments yielded 94 and 93% of their fruit with a diameter greater than 70 mm at harvest (Supplementary Fig. S1). Fruit FW and volume also increased with reduced crop load and competition for photosynthates, with a two-fold increase in both FW and volume from the unthinned when compared to the extreme thinning (30 cm) treatment (Table 1; Fig. 1). Average fruit FW only exceeded 200 g with the 15 cm and 30 cm thinning treatments (Table 1). Return bloom and fruit set increased with reduced carbohydrate competition during flower bud initiation in the previous season amongst all thinning severities, when compared to the unthinned control (Table 1). However, return fruit set was maximized with the 15 cm thinning treatment (Table 1). Minimal impacts were seen in the leaf elemental content across crop loads. Specifically, significant differences were observed only for K, with elevated levels in leaves from the 15 cm and 30 cm spacing treatments compared to the unthinned control and other treatments (Supplementary Table S1).

Non-destructive simultaneous internal quality (DMC, SSC) and maturity (I_AD_) analyses (Minas *et al*., 2021) using an accurately calibrated (high *R*^*2*^ values and low RMSEP in Fig. 2A, B, C) handheld NIRS sensor for ‘Cresthaven’ peach (Minas *et al*., unpublished) demonstrated significant differences across all parameters assessed (Table 2). Dry matter content and SSC increased significantly by 25 and 27%, respectively, from the heaviest to the lowest crop load (corresponding to a decrease in carbon competition) (Table 2; Fig. 2D, E). However, the degree of differences diminished between the 10 and 30 cm fruit spacing treatments (Table 2). Interestingly, 99 and 98% of fruit from the 15 and 30 cm fruit spacing exceeded a DMC level of 11%, while only 31% of fruit surpassed this level in the unthinned treatment (Fig. 2D). Similarly, fruit from same thinning treatments had 87% of its fruit surpassing an SSC level of 11%, a minimum consumer quality standard (Hilaire, 2003), while the unthinned treatment had only 8% of its fruit surpassing this threshold (Fig. 2E). Maturity (I_AD_) advanced (lower I_AD_ values) with increased thinning severity, with fruit that were coming from unthinned and heavily cropped (5 cm fruit spacing) trees being less ripe than the fruit coming from the remaining thinning (10, 15 and 30 cm) treatments at harvest (Table 2). At harvest, 69% of fruit were classified as immature (> 0.60 I_AD_) in the unthinned control, while the 15 cm treatment gave less immature fruit (57% > 0.60 I_AD_) (Fig. 2F). Flesh firmness (FF) exhibited a weak relationship between the NIRS estimation and actual destructive values (Minas *et al*., 2021; Minas *et al*., unpublished), so only the destructive values are presented (Table 2). Minimal differences were detected in FF across the five crop load treatments, with only fruit from the 15 cm treatment being significantly softer than fruit from the 5 cm treatment (Table 2). Overall, these data demonstrate that increased thinning severity (corresponding to increased carbon availability) resulted in improved fruit quality characteristics and advanced maturity at harvest. However, it is important to note that these data do not confirm if these observed quality shifts are direct results of carbon availability due to the differing fruit-to-fruit competition and/or the advancement/delay in fruit maturation.

**Fig. 2.**
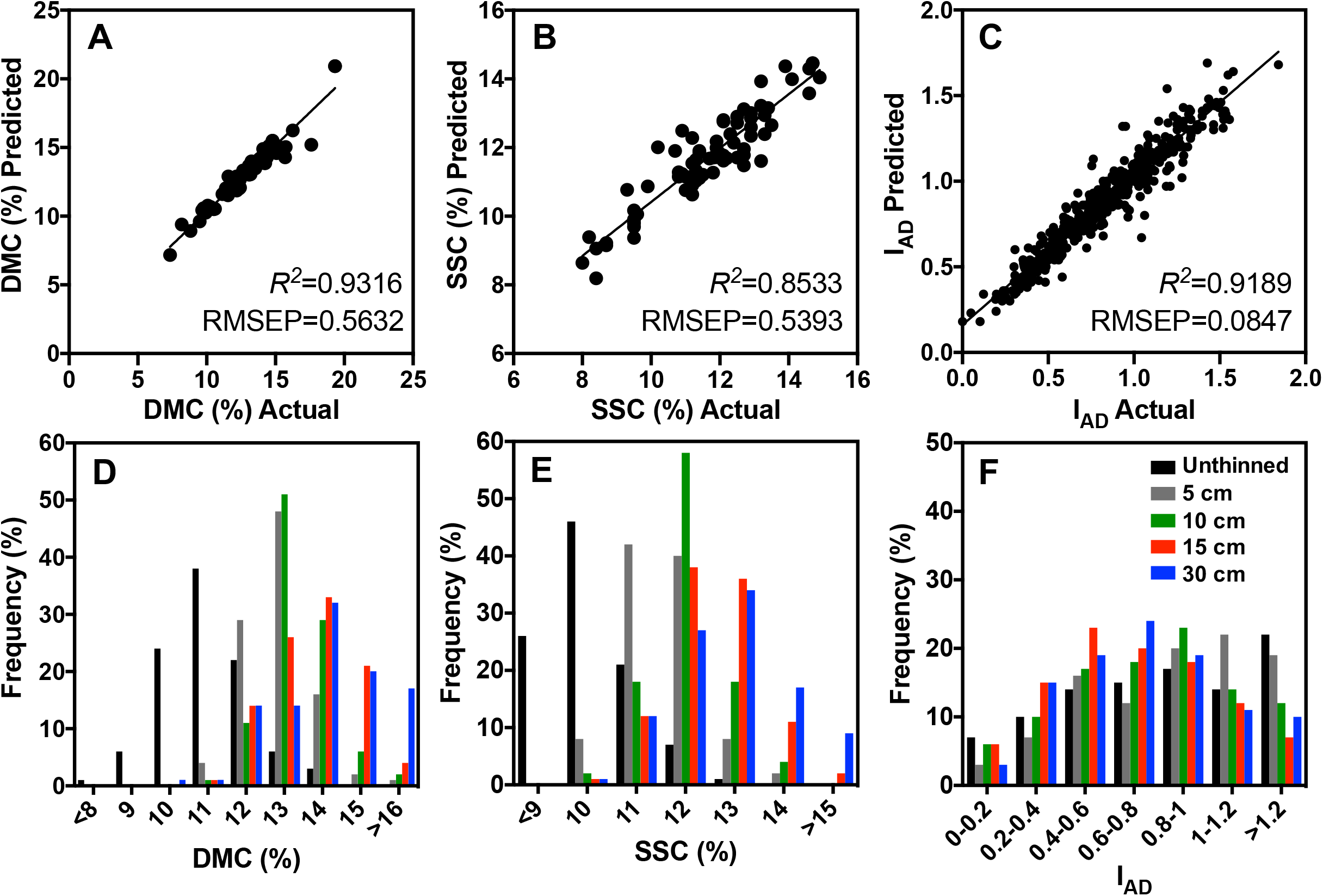
Non-destructive quality model validation and distribution of quality parameters at harvest across variable carbon supply conditions. Multivariate regression models for non-destructive estimations of peach internal fruit quality and maturity were validated with fruit samples at harvest (138 DAFB) from the various carbon supply treatments. Non-destructive predicted estimations were plotted against actual destructive values for validation of quality and maturity parameters: dry matter content (DMC) (A), soluble solids concentration (SSC) (B) and index of absorbance difference (I_AD_) (C). The created models for these parameters were evaluated for linearity (*R*^*2*^) and root mean square error of prediction (RMSEP) to demonstrate the accuracy of the model in estimating maturity and various quality parameters with near-infrared (NIR) spectroscopy (A-C). The impact of thinning severity on ‘Cresthaven’ fruit distributions across classes of DMC, SSC and I_AD_ are shown (D-F) and were assesses using the non-destructive NIRS multivariate models. Various thinning treatments’ frequencies for each parameter are visualized as: unthinned (black), 5 (grey), 10 (green), 15 (red) and 30 cm (blue) (D-F).

### Phenotyping fruit of equal maturity demonstrates superior quality in thinned fruit, especially at S4

In order to determine the true impact of carbon supply on fruit quality development, fruit of equal maturity from one thinning treatment (15 cm, carbon sufficient) were compared with the unthinned (carbon starved) control (Fig. 1E, F and 3) during peach growth and development. As described above, a handheld non-destructive NIRS sensor was utilized to sort for fruit of equal maturity at each developmental stage (S2, S3 and S4) between the carbon starved (unthinned) control and carbon sufficient (thinned at 15 cm fruit spacing referred from this point forward as “thinned”) treatments (Fig. 3F). Sorting for equal maturity allows for comparisons of preharvest treatments without the confounding influence of variable maturity. Maturity control was confirmed by ensuring that I_AD_ values did not differ between the crop load treatments at each developmental stage (Fig. 3F). Selected fruit from the thinned treatment and the unthinned control exhibited I_AD_ values that decreased by 66 and 63% over time from S2 to S4, respectively (Fig. 3F). In addition, FF decreased relatively similarly across the developmental stages, with an 81 and 73% reduction in firmness from S2 to S4 in the thinned treatment and unthinned control, respectively (Fig. 3E).

**Fig. 3.**
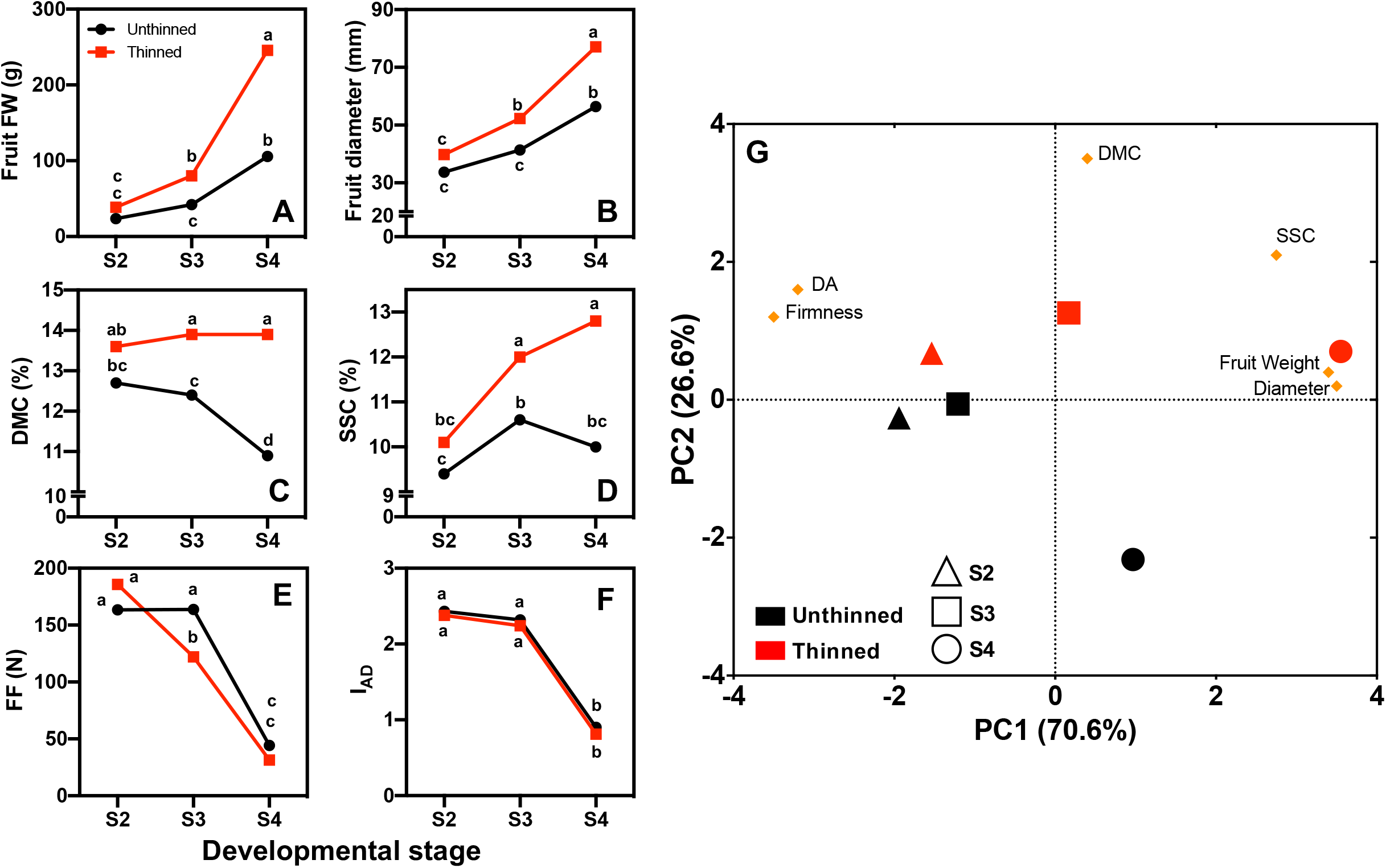
The true impact of carbon supply on fruit internal quality development. Impact of two carbon supply treatments: unthinned (starved) and thinned (15 cm, sufficient) on fruit fresh weight (FW, g; A), diameter (mm; B), dry matter content (DMC, %; C), soluble solids concentration (SSC, %; D), flesh firmness (FF, N; E), and physiological maturity (I_AD_; F) from destructive quality analysis on 10 fruit per treatment at each developmental stage. Means of fruit plotted at three developmental stages: stage 2 (S2, pit hardening, 72 DAFB), S3 (cell elongation, 109 DAFB), and S4 (harvest, 138 DAFB). Error bars are not visualized as they were too small to be seen in the figure. Fruit quality was assessed on fruit of equal maturity, assessed with I_AD_, at each developmental stage (F). Two-way ANOVA (thinning treatment x developmental stage) was used to detect differences across means with lettering assigned by Tukey’s HSD test. Means with the same letter indicate non-significance at *P*<0.05. Principal component analysis (PCA) was also conducted to assess the impact of variable carbon supply on peach fruit phenotype. Thinning treatment (red) vs. unthinned fruit (black) and developmental stage [S2 (triangle), S3 (square), S4 (circle)] scores are scaled on the PCA with peach fruit quality parameters (loadings, orange diamonds). Large symbols on the PCA are the averaged ten fruit per each thinning x developmental stage treatment. The PCA demonstrates that developmental stage was a major contributor for quality variation as indicated by separation on PC 1 (∼ 71%). Additional quality variation due to the thinning treatment and other factors are visualized on PC 2 (∼ 27%). PCA shows minimal difference in fruit phenotype at S2, while at S4, phenotypes between thinning treatments (Fig. 1E) are highly separated.

In general, fruit quality parameters were not significantly different between the thinned treatment and unthinned control at S2, but improved fruit quality in the thinned treatment was observed at S3 and S4 (Figs. 1D, E and 3A, B, C, D). For example, fruit FW and diameter increased faster and to heavier/larger levels (1.5-fold heavier and 37% larger) at harvest (S4) than the unthinned control (Figs. 1D, E and 3A, B). For both fruit FW and diameter, fruit from the unthinned control at S4 were equivalent to fruit from the thinned treatment at S3 (Fig. 3A, B). Similar trends were observed for DMC and SSC. Specifically, a 28% increase in both DMC and SSC were observed in fruit from the thinned treatment when compared to the unthinned control at S4 in fruit of equal maturity (Fig. 3C, D). Importantly, both of these carbohydrate-based quality parameters (DMC and SSC) increased and remained higher in the fruit from the thinned treatment representing a condition of low fruit-to-fruit competition. Conversely, in the unthinned control, these parameters decreased as fruit reached commercial harvest maturity (between S3 and S4), reflecting the magnitude of the carbon starvation condition (Fig. 3C, D). These results suggest that the thinning treatment improved carbon supply to the fruit, resulting in enhanced fruit quality.

Global changes in fruit quality can be visualized using principal component analysis (PCA, Fig. 3G) further demonstrating the trends described above. At S2 (triangles, Fig. 3G) fruit from the thinned treatment and unthinned controls are highly similar (Fig. 1E). However, at S3 and S4 (squares and circles, Fig. 3G) it is clear that the overall fruit quality between the thinned treatment and the unthinned control diverges (Fig. 1E). Evaluation of the influence of the loadings (presented as orange diamonds in the PCA biplot in Fig. 3G) indicates increased values in quality parameters (DMC, SSC, FW, and fruit diameter) in fruit from the thinned treatment at S3 and S4.

### Elemental composition in peach fruit mesocarp is affected by maturation and early carbon sufficiency/starvation

Ionomic analysis was conducted to evaluate the impact of carbon limitation/availability across developmental stages on elemental composition of equal maturity peach mesocarp. The analysis screened for a panel of 23 elements, of which, 3 (Li, Pb, and W) were below the assay LOQ (Supplementary Table S3). Principal component analysis demonstrates a large percentage of the variation being attributed to PC1 (66.6%) representing developmental stage, with an additional portion of the variation explained with PC2 (13.1%) representing carbon supply and likely other unknown factors (Supplementary Fig. S2). The concentration of detected elements decreased in both treatments with the advancement of peach fruit growth, development and maturation (Supplementary Fig. S3). At S2, 10 elements (Al, As, Ba, Ca, Cd, Fe, K, Mg, Sr, V and Zn) had significantly higher concentrations in the unthinned control compared to the thinned treatment (Supplementary Table S3). At harvest (S4), only 4 elements (Al, As, Sr and V) remained significantly higher in the unthinned controls (Supplementary Table S3). Taken together, these results demonstrate that there is an overall trend in decreasing elemental concentration with developmental stage advancement, and that most of the significant differences due to the carbon availability that were observed early in development, were no longer relevant at harvest.

### Vast metabolic differences shift early on during fruit growth and development as a response to carbon sufficiency/starvation

In total, 189 metabolites were detected in peach mesocarp samples (Fig. 4). Of those, 36 metabolites were confidently annotated (orange diamonds, Fig. 4). Principal component analysis revealed that a large percentage of the metabolic variation is attributed to PC1 (34.1%) representing fruit developmental stage, with additional variation explained by PC2 (14.7%) representing carbon supply manipulation (Fig. 4). Importantly, the largest metabolic separation was observed between the thinned treatment (red symbols) and unthinned control (black symbols) at S2 (triangles). This separation is reduced at S3 (squares) and further at S4 (circles). These early shifts in the metabolome resulting from variable carbon supply levels due to thinning treatment are in direct contrast to the observed trends in fruit phenotype, which were greatest at the last developmental stage (Figs. 1D and 3). In other words, this early dramatic shift in the metabolome (Fig. 4) is in direct contrast to phenotypic shifts, where PCA separation at S2 in Fig. 3G is minimal. Overall, shifts in the metabolome (Fig. 4) appear to behave inversely with shifts in fruit quality and maturity attributes (Fig. 3G).

**Fig. 4.**
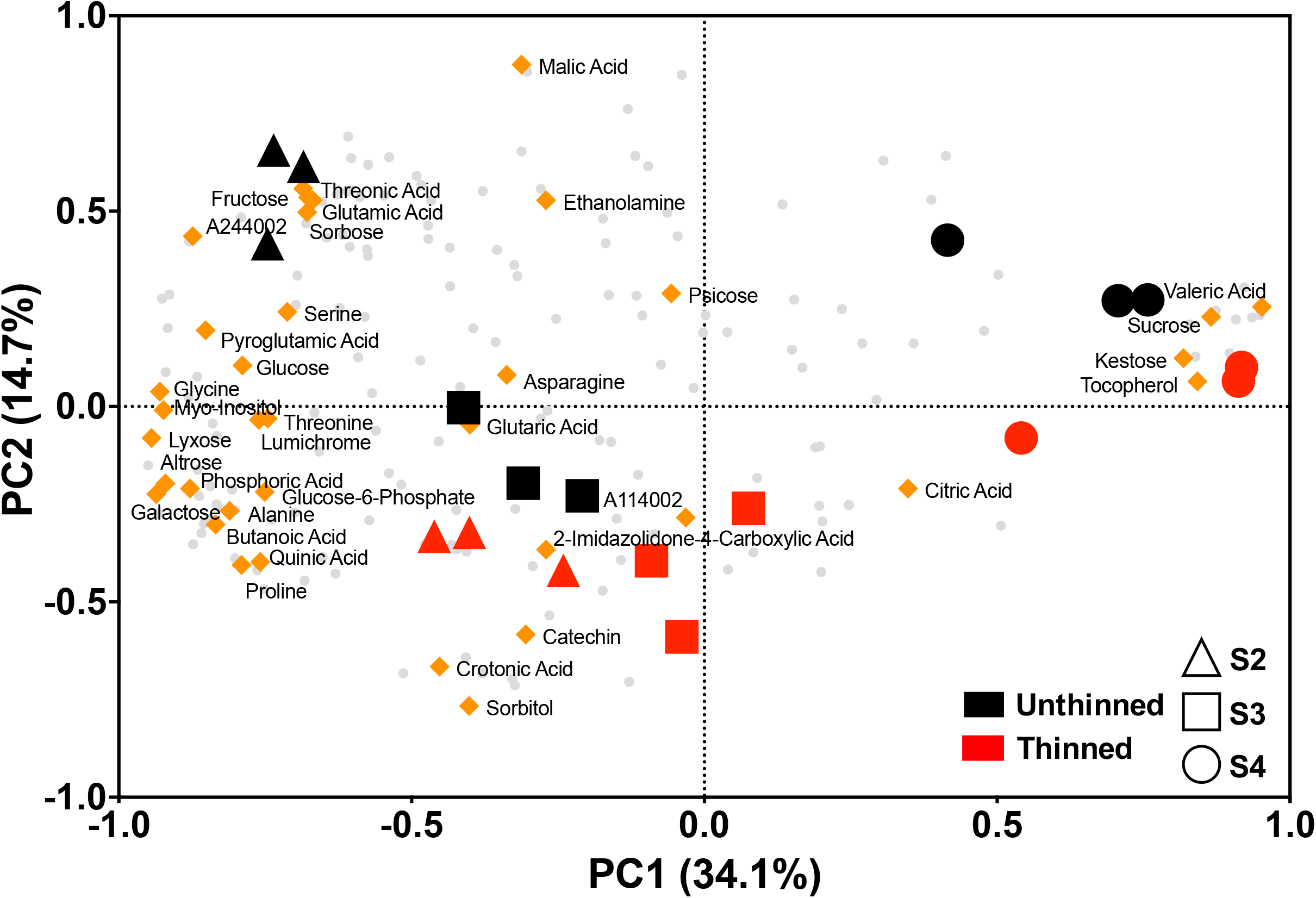
Principal component analysis biplot of variable carbon supply on peach fruit metabolism. Large symbols indicate the scores for the thinning treatments [unthinned (black) vs. thinned (red)] and developmental stages [S2 (triangle), S3 (square), S4 (circle)] and are scaled with the metabolites detected in the peach mesocarp (loadings). Principal component analysis (PCA) of the three reps per each thinning x developmental stage treatment demonstrate that developmental stage was a major contributor for metabolome variation as indicated by separation on PC 1 (∼ 34%). Additional metabolome variation due to the carbon availability and other factors is visualized on PC 2 (∼ 15%). The 36 annotated metabolites are shown on the PC loadings as orange diamonds. The remaining 152 peaks that were detected, but not annotated, are shown as grey circle loadings in the background. PCA shows wide variation in metabolome between carbon supply treatments at S2, while at S4, fruit metabolome scores are minimally different.

General shifts in the peach metabolome across developmental stages include a decreasing abundance of various amino and organic acids (Fig. 5). Some hexoses decreased with fruit development (e.g. glucose, fructose and galactose), while more complex saccharides increased with development (e.g. sucrose and kestose; Fig. 6). Additional metabolites showing this trend include valeric acid and tocopherol (Fig. 6). Metabolites including myo-inositol, proline and butanoic acid showed the inverse trend, decreasing significantly throughout development (Figs. 5, 6, Supplementary S4). Fructose, sorbose, myo-inositol and threonic acid were all significantly more abundant in the carbon starved, unthinned control at S2, by 75, 73, 79 and 67%, respectively (Figs. 6, Supplementary Fig. S5). Interestingly, these differences became less significant as fruit development progressed (Fig. 6). Conversely, sorbitol, quinic acid and tocopherol were significantly more abundant by 46, 50 and 47%, respectively, in the thinned treatment at S2 (Fig. 6).

**Fig. 5.**
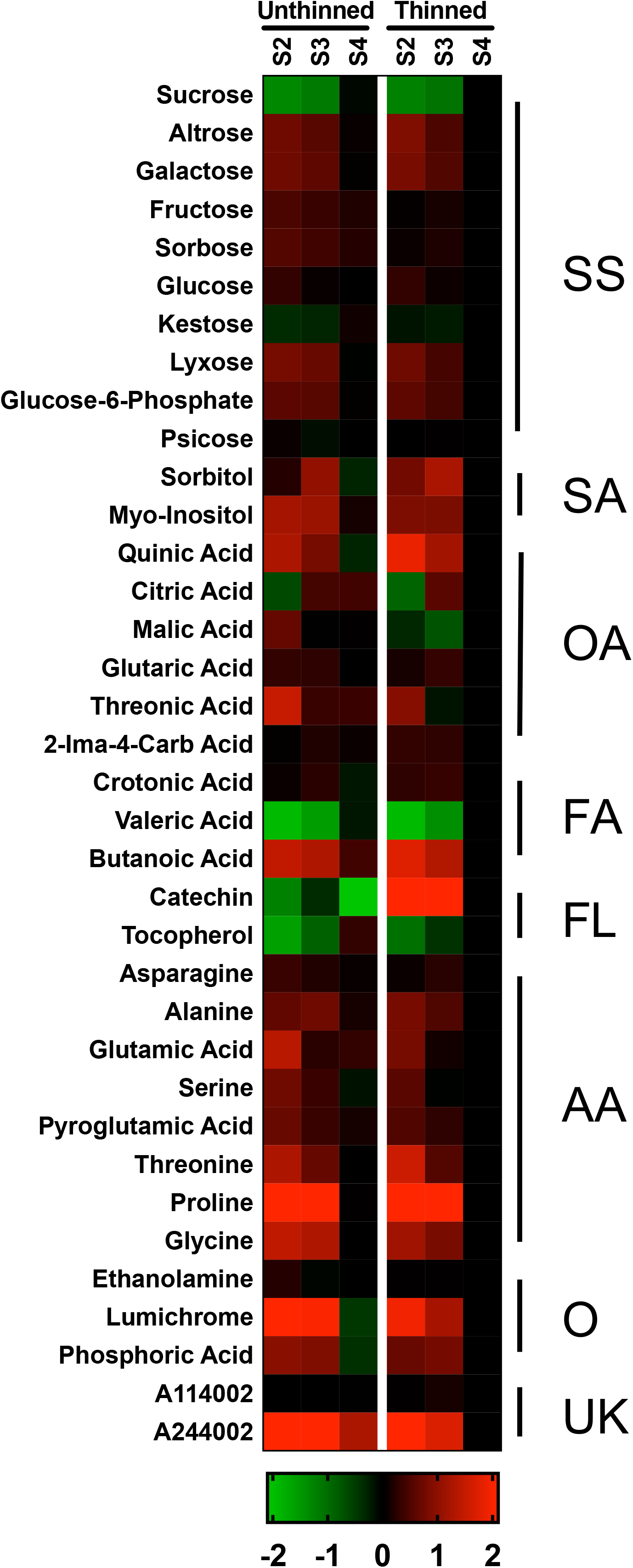
Heat map of metabolite profiles across development under two carbon supply conditions. Profiles of metabolism changes across two carbon supply conditions during growth and development in peach fruit mesocarp. Figure shows comparisons of the metabolite abundance across S2, S3 and S4 in the carbon starved (unthinned) and sufficiency (thinned) fruits. The ratio between thinning x developmental stage treatments were compared to the “optimal” conditions of thinned-S4 (far right column). Each of the 36 annotated metabolites were transformed into log_2_ and shown with the following color scale (green to red) according to Lombardo *et al*., (2011). Fruits at each developmental stage were of equal maturity according to the I_AD_ measured by the DA meter (Fig. 3F). Annotated metabolites are organized by chemical class: soluble sugars (SS), sugar alcohols (SA), organic acids (OA), fatty acids (FA), flavonoids (FL), amino acids (AA), other (O) and classified unknowns (UK).

**Figure 6.**
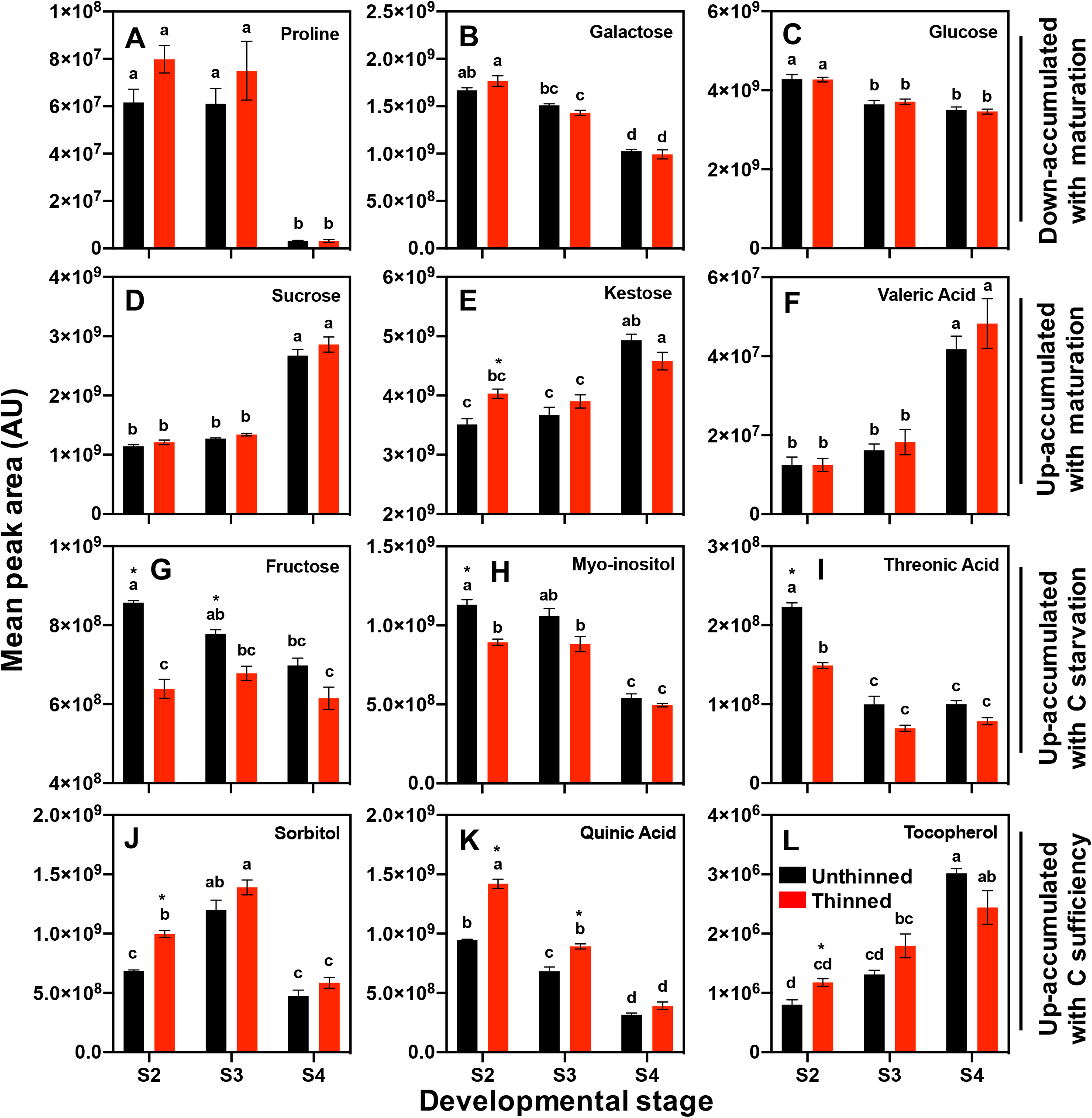
Accumulation trends of metabolite abundances throughout fruit maturation and across variable carbon supply treatments. Figure showcases the mean peak area (AU) of selected metabolites that are down-accumulating (A-C) or up-accumulating (D-F) throughout peach fruit development and maturation (S2 – S4) in two carbon supply conditions. The mean peak area of selected metabolites that are up-accumulating with carbon starvation, e.g. decrease in thinned vs. unthinned (G-I) or with carbon sufficiency, e.g. increased in thinned vs. unthinned (J-L) are also shown across development. The bars indicate carbon supply treatments: unthinned (black) and thinned (red, 15 cm fruit spacing). Samples were controlled for equal maturity (I_AD_) across development and between carbon supply treatments. Mean values ± S.E. are displayed. Means followed by the same letter are not statistically different according to Tukey’s HSD test (*P*<0.05). * indicates a significant difference between carbon supply treatments at a singular developmental stage according to a one-way ANOVA (*P*<0.05).

### Metabolite accumulation and degradation throughout development correlate with fruit physiological attributes

Of the 36 annotated metabolites, 25 strongly correlated (*R*^*2*^>0.70 or <-0.70) with at least one fruit physiological characteristic (fruit FW, fruit diameter, DMC, SSC, FF and I_AD_) throughout development in fruit from the thinned treatment (15 cm fruit spacing; Fig. 7), which is a commonly used commercial thinning practice. Sucrose demonstrated extremely strong positive correlations with fruit FW (*R*^*2*^=0.97), diameter (*R*^*2*^=0.95) and SSC (*R*^*2*^=0.71) (Fig. 7). Kestose (saccharide), along with tocopherol (i.e. antioxidant; vitamin E) exhibited a strong positive correlation with fruit FW (*R*^*2*^=0.74 and 0.79, respectively). Glucose-6-phosphate was the only metabolite to have a strong correlation (*R*^*2*^<-0.70) with DMC across fruit development (Fig. 7). Many strong negative correlations were observed for SSC including several monosaccharide hexoses (lyxose, altrose, galactose, and glucose; Fig. 7). Other metabolites showing strong negative correlations with SSC include organic and fatty acids such as glutamic acid, butanoic acid, threonic acid, pyroglutamic acid, and quinic acid (*R*^*2*^=-0.90). Many positive correlations between physiological maturity and metabolite abundance were also observed. For example, as I_AD_ values decreased (maturity advancement), the abundance of several amino acids and monosaccharide hexoses (e.g. glucose, fructose and galactose) decreased, while the abundance of di- and polysaccharides increased (e.g. sucrose and kestose; Figs. 5, 6 and 7). Strong positive correlations with I_AD_ was observed for multiple amino acids including alanine, glycine, and proline (*R*^*2*^=0.93). A strong negative correlation with I_AD_ was observed for sucrose (*R*^*2*^=-0.97), however positive correlations were observed for other monosaccharides including glucose, altrose, galactose (*R*^*2*^=0.89). The strongest metabolite-metabolite relationships (positive/negative) include altrose with quinic acid (*R*^*2*^=0.99) and sorbitol with malic acid (*R*^*2*^=-0.95).

**Fig. 7.**
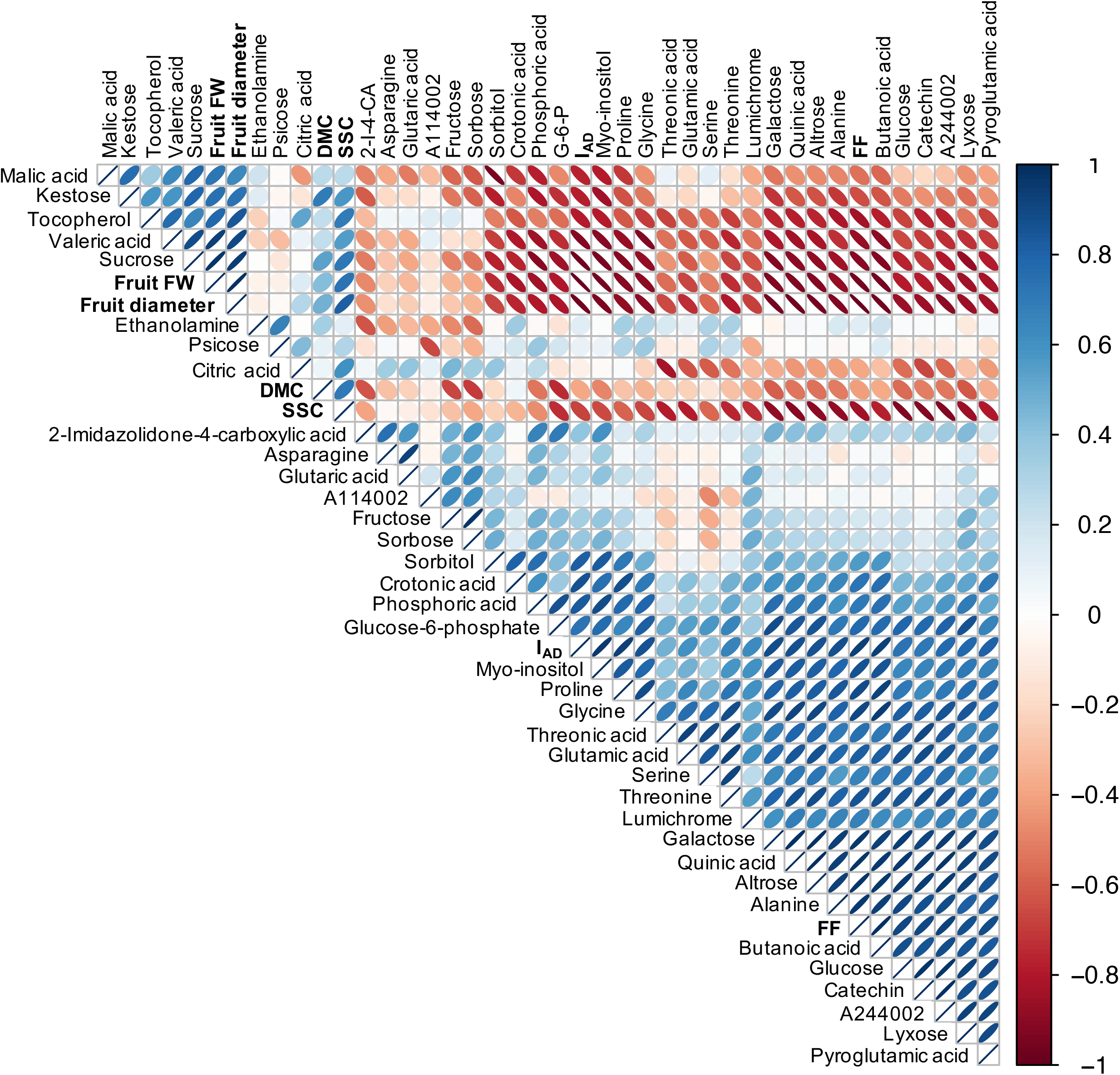
Heat map of correlation matrix between fruit physiological characteristics and metabolite abundances. Heat map for 42 physiological (6) and metabolic (36) characteristics was created based on hierarchical clustering on the spearman *R*^*2*^ values. Colors and ellipse eccentricity visualize the direction and strength of the relationship between characteristics. Data used for relationships are from the carbon sufficient (thinned, 15 cm) treatment only, across all developmental stages (S2, S3 and S4). Correlations show the relationship between the accumulation or degradation of fruit physiological parameters and metabolite abundances throughout maturation.

### Catechin and a classified unknown metabolite represent different relationships with fruit quality parameters at harvest

At the S2 developmental stage, 16 of the 36 annotated metabolites were significantly different between the unthinned control (carbon starvation), and the sufficiently carbon supplied, thinned treatment (*P*<0.05; Supplementary Table S4). Whereas only two of these metabolites remained significantly different at S4 (Supplementary Table S4), catechin and a classified unknown, A244002 (Figs. 5, 8A, D). Catechin remained highly elevated in the thinned treatment throughout development, but the magnitude was greatest at S2 (47-fold; Fig. 8A). At S3 and S4, catechin was still higher in the carbon sufficient condition when compared to the starved, but to a lesser degree (7-fold and 4-fold, respectively; Fig. 8A). The opposite trend was observed for A244002, with a greater abundance (2-fold) observed throughout fruit development in the carbon starved condition compared to carbon sufficiency (Fig. 8D).

**Fig. 8.**
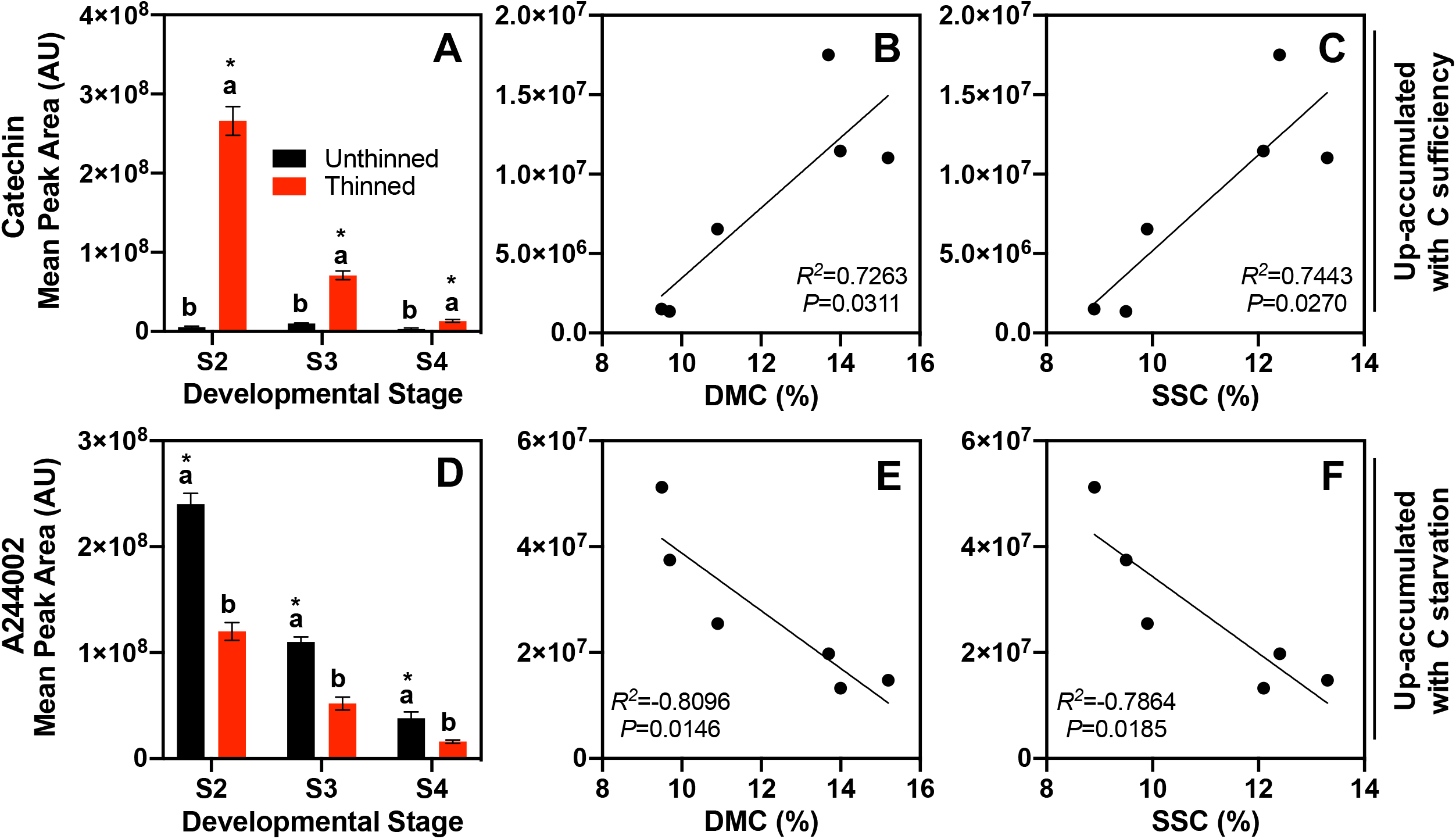
Abundance of proposed metabolite primers and their relationship with peach internal quality parameters at harvest. Mean peak area (AU) of catechin (A) and of a classified unknown (A244002, D), respectively, at each developmental stage (S2, S3 and S4) between carbon sufficiency treatments [unthinned vs. thinned (15 cm fruit spacing)]. Mean values ± S.E. are displayed. Means followed by the same letter are not statistically different according to Tukey’s HSD test (*P*<0.05). * indicates a significant difference between carbon supply treatments at a singular developmental stage according to a one-way ANOVA (*P*<0.05). The relationships between the mean peak area of catechin and A244002 with dry matter content (DMC, %; B and E, respectively) and soluble solids concentration (SSC, %; C and F, respectively) at harvest (S4) with three replicated samples from both the carbon availability treatments are plotted. *R*^2^ values are displayed to demonstrate the linearity of the relationships, along with *P*-values to indicate the significance of the relationship.

At harvest (S4), these two compounds were assessed for their relationship with the physiological internal quality parameters related to photosynthates (SSC and DMC), in both the carbon sufficient (thinned) and the carbon starved (unthinned control. Catechin exhibits positive and significant (*P*<0.05) correlations at harvest with both DMC (*R*^*2*^=0.73) and SSC (*R*^*2*^=0.74), indicating superior fruit quality (Fig. 8B, C). Alternatively, A244002 showed strong negative and significant correlations (*P*<0.05) with DMC (*R*^*2*^=-0.81) and SSC (*R*^*2*^=-0.79) at harvest, signaling inferior quality (Fig. 8E, F).

## Discussion

### Increased thinning severity enhances fruit quality, but is it a result of crop load management or advanced maturation?

Reducing the sink number on trees (via fruit thinning) allows for increased photosynthates for the remaining fruit, which enhances quality and advances maturity (Minas *et al*., 2021). Determining an optimal crop load for a particular cultivar in a growing region must balance fruit production and quality standards. In our study, increased thinning severity reduced yields, but enhanced fruit quality and advanced fruit maturity (Tables 1 and 2). Among the different crop load management strategies studied the 15 cm fruit spacing (2.6 fruit · cm^-2^ of TCSA) provided an acceptable yield (∼40 kg tree^-1^ or 20 MT · ha^-1^) for mid-density late-ripening peaches in Colorado growing conditions (USDA, 2019) with commercially acceptable fruit FW, size and quality to secure grower profitability (Minas *et al*., 2018). At the same time, 15 cm fruit spacing maximized return fruit set and had higher return bloom numbers when compared to the rest of the crop load treatments (Table 1). This demonstrates evidence that a sufficient carbon supply was achieved, allowing for optimum fruit quality development, balanced vegetative growth (Grossman and DeJong, 1995) and adequate floral bud differentiation and fruit set for the next season. Collectively, these production, quality and reproductive capacity indicators make 2.6 fruit · cm^-2^ of TCSA (i.e. 15 cm spacing in this experiment) a potential target for consistent high yields of high-quality late-ripening peach fruit.

This trade-off of yield for quality has been documented in several peach crop load studies, where reduced crop loads improve internal fruit quality (Inglese *et al*., 2002; Marini, 2003; Alcobendas *et al*., 2012). However, the reported quality enhancements may be unintentionally confounded by maturity advancement in the lower crop loads and thus not truly linked to the preharvest factor alone (Minas *et al*., 2018). Therefore, our study sought to evaluate fruit of equal maturity, sorted by a handheld NIRS sensor, to understand the actual impact of thinning on fruit internal quality development. With the selection of the 15 cm fruit spacing thinning treatment as the optimal crop load management strategy, comparisons were made between this treatment and the unthinned control for fruit quality parameters, assessed at equal maturity, as confirmed with I_AD_ (Fig. 3F).

At S2, there were no differences in any physiological parameter between the thinned treatment and unthinned control (Fig. 3A-F), while the phenotype of these fruits at this stage were very similar (Fig. 1E). However, initiating at S3, both phenotype and fruit quality characteristics begin to diverge (Figs. 1E and 3G). At S4, fruit phenotypes between the thinned treatment and unthinned control were drastically different, in respect to size, weight and overcolor (Figs. 1D, E, 3A-F, Supplementary Fig. S1, and Table 1). Dry matter content and SSC increased throughout development in thinned fruit but begin to decrease between S3-S4 in unthinned control (Fig. 3C-D). A high number of fruit during the critical stages of fruit growth and development, could generate a sink demand that can be too high and fruits can become source (carbon) limited (Grossman and DeJong, 1995). This leads to deleterious effects on fruit quality in the unthinned trees as a result of diluted photosynthate availability. Multiple studies have demonstrated that increased crop load reduces DMC and SSC accumulation (Berman and DeJong, 1996; Crisosto *et al*., 1997; Cirilli *et al*., 2016; Minas *et al*., 2021), however these may be attributed to maturation delay. Here, by controlling for fruit of equal maturity we demonstrate that these reductions in internal fruit quality are directly related to crop load and carbon availability during fruit growth and development.

### Ionome variation linked mainly with peach fruit development and maturation, and less with carbon availability

Overall, the ionome appears to shift with peach fruit development and maturation (Supplementary Fig. S2). The concentrations of nearly all elements decreased over time, with little variation in elemental composition between the thinned treatment and unthinned control (Supplementary Fig. S3). Elemental differences are more abundant in S2 than at harvest (S4, Supplementary Table S3). Specifically, at S2, 10 elements were statistically different (*P*<0.05), while only 4 elements were different at harvest (S4) (increased concentrations of Al, As, Sr and V in the unthinned control) (Supplementary Table S3). All 10 elemental differences at S2 indicate higher concentrations in the unthinned treatment as well, including key nutrients such as Ca, Fe, K, Mg, and Zn. This may be a stress response of the tree, to flood the fruitlets early with critical nutrients to support the heavy fruit load due to high competition conditions or to reduce fruit expansion rates (Wünsche and Ferguson, 2005). However, as the tree’s nutrients get exhausted, or as nutritional support may be more related to maturation, less differences in elemental composition appear. Previous studies in apple have observed more significant variation in elemental composition with crop load treatment (Wünsche and Ferguson, 2005), however, this result is likely confounded by fruit maturity. The results presented here from fruit of equal maturity suggest that carbon availability has minimal impact on elemental composition at harvest.

### Peach fruit metabolites are related to fruit growth, development and maturation processes

Fleshy fruit development, maturation and ripening are intricately regulated by physiological and molecular processes (Giovannoni *et al*., 2017). Therefore, to understand any preharvest factor effect on peach fruit metabolism, samples must be evaluated at equal maturity to eliminate this confounding variable (Minas *et al*., 2018). The novel methodological approach of this study enabled the opportunity to investigate the true impact of carbon manipulation on peach fruit metabolism and to generate a baseline understanding of metabolites that are involved in fruit growth and development. Importantly, most of the metabolome variation observed is reflective of fruit developmental changes rather than carbon manipulation (Figs. 4, 5). Thus, our data suggest that metabolic shifts are intricately connected to the highly coordinated processes of fruit maturation and ripening (Giovannoni *et al*., 2017).

Since maturity control was validated using I_AD_ (Fig. 3F), abundances of specific compounds could be linked with developmental stage and maturation. Previously, it has been suggested that physiological maturity could be determined metabolically based on the abundances of sucrose and/or quinic acid in peach fruit mesocarp and seed (Chapman *et al*., 1991). Sucrose continues to accumulate throughout peach development, reaching maximum levels at harvest, while quinic acid behaves inversely, starting high and depreciating throughout maturation (Chapman *et al*., 1991; Wu *et al*., 2005). Our data confirm these trends and reveal no differences in sucrose nor quinic acid across carbon supply treatments at harvest (S4; Fig. 6D, K). Furthermore, there are minimal differences (only two compounds) in the metabolome between carbon sufficiency and carbon starvation treatments at S4 (Fig. 5 and Supplementary Table S4). This result underscores the strong connection between the primary metabolism and fruit growth and development, as these compounds are fundamental for essential fruit functions. In other words, the lack of observed differences in metabolites reflective of the primary metabolism between carbon sufficiency/starvation conditions at harvest, is likely a result of controlling for maturity.

Additional metabolites that strongly correlate with fruit maturation are presented in Fig. 6. Similar to previous research, our results show that glucose, fructose and galactose, along with various organic acids, are highly abundant in immature fruit and decrease in abundance throughout peach fruit development, while sucrose steadily increases in abundance until ripening, where it rapidly increases and becomes the most abundant saccharide at harvest (Fig. 6; Chapman *et al*., 1991; Wu *et al*., 2005; Lombardo *et al*., 2011; Bae *et al*., 2014). These saccharide conversions from mono-to polysaccharides across maturation are supported in the literature (Cirilli *et al*., 2016), which demonstrates the conversion of simple hexoses into more complex sugars throughout development (Figs. 5, 6). Sorbitol has also been linked to fruit development, showing a peak in sorbitol abundance at the S2/S3 transition followed by a decline towards S4, which is supported with our results (Cirili *et al*., 2016; Figs. 5, 6).

Several monosaccharides and sugar alcohols detected in this study, under sufficient carbon supply (thinned treatment), show inverse correlations with FW and SSC and decreased with maturation (Fig. 7). For example, galactose and glucose abundances are positively correlated with I_AD_ (*R*^*2*^=0.89 and 0.73, respectively) and firmness (*R*^*2*^=0.96 and 0.91, respectively), meaning their abundances decrease with maturity advancement and flesh softening. Textural changes and flesh softening in pome and stone fruit has been attributed to the solubilization of cell wall components and pectic substances, such as saccharides (Nara *et al*., 2001; Brummell *et al*., 2004). Galactose, found in cell walls, has been documented to decrease with flesh softening, as it is solubilized throughout maturation due to the activation of polygalacturonase (Brummell *et al*., 2004). Complex carbon compounds, such as sucrose and kestose, demonstrate opposite trends, increasing with maturation (*R*^*2*^=-0.97 and -0.78, respectively; Fig. 6), FW (*R*^*2*^=0.97 and 0.74, respectively) and SSC (*R*^*2*^=0.71 and 0.58, respectively; Fig. 7).

The most significant changes observed for amino acids across maturation were for proline, which was not significantly impacted by carbon availability (Fig. 7). Proline during fruit growth and development was also highly positively correlated with I_AD_ (*R*^*2*^=0.93). Other amino acids including glycine, threonine and alanine were positively correlated to firmness and maturity (I_AD_), and negatively correlated with weight and SSC (Fig. 7). These results are consistent with previous studies showing that amino acids are typically elevated in immature fruit/early and decrease throughout development (i.e. when firmness and I_AD_ values drop, and weight and SSC increase) (Monti *et al*., 2016).

Malic acid showed a positive correlation with FW and a negative correlation with maturity advancement (i.e. I_AD_ decrease) (Figs. 6, 7). This result supports the use of malic acid abundance as a maturity index and it has also been shown to correlate well with consumer preference (Chapman *et al*., 1991; Colaric *et al*., 2005). Organic acids typically accumulate in immature fruit and decrease with fruit development (Bae *et al*., 2014). Valeric acid, a fatty acid, does not follow this general trend, reaching a maximum abundance in S4. This suggests that it may be involved in fruit ripening and aroma volatilization. Butanoic acid, another fatty acid, which has been shown to be a predominant aromatic compound associated with the onset of ripening in banana (Zhu *et al*., 2018), was found to decrease with peach maturation (Supplementary Fig. S4).

### Early sufficient carbon supply allows for metabolic investments that are priming fruit quality potential

There are apparent differences in the metabolite profiles between carbon supply treatments, although this variation appears be more significant early on, immediately after thinning in S2 (Figs. 4, 5). Upon S4, few compounds were significantly different between the carbon sufficient (thinned) and starved treatments (unthinned) (Supplementary Table S4, Figs. 6, 8A, D), underscoring the high level of metabolite regulation throughout maturation and justifying our methodology of sampling fruit of equal maturity. It is significant to note that at S2, there is a high level of metabolic variation, while there are very few differences in quality characteristics and phenotype (Figs. 1, 3-5). At S4, most metabolic differences appear to dissipate, while there are extreme differences expressed in quality and phenotypic characteristics (Figs. 1, 3-5). The extreme differences in inter-fruit carbon competition conditions appears to create an early and dramatic metabolic shift that primes quality development.

With sufficient carbon supply (adequate fruit thinning), the priming appears to lead to quality enhancement, while the carbon stressed unthinned control (high carbon competition), leads to quality detriment. The importance of early action for crop load management to maximize carbon resources for the developing fruit is strongly recognized (Grossman and DeJong 1995). This is because under sufficient carbon supply (via thinning), fruit can achieve their maximum fruit growth potential (Grossman and DeJong, 1995). Similarly, our results further suggest that this early carbon supply adjustment initiates vast metabolic changes in the developing fruits, which allows them to achieve maximum fruit quality potential as well.

Fruit from unthinned trees had a high abundance of several organic acids at S2, such as citric acid, malic acid and threonic acid (Supplementary Table S4), which have been previously correlated with a lack of sweetness in peach fruit (Colaric *et al*., 2005). Furthermore, threonic acid had a negative correlation with SSC in the carbon sufficient treatment throughout development (*R*^*2*^=-0.79), which also supports the lack of quality at harvest in the unthinned controls (e.g. reduced SSC and DMC) (Figs. 3, 6 and 7). Additionally, the accumulation of particular organic acids has been shown to be a response to abiotic stresses (e.g. salt, soil acidity, drought) in other species (Timpa *et al*., 1986; Fougere *et al*., 1991; Zeng *et al*., 2008). Thus, our results suggest that carbon limitation, resulting from a high crop load (i.e. unthinned control), may induce a similar abiotic stress response, as reflected with an elevated abundance of organic acids at S2 (Table S4).

Multiple carbohydrate-related metabolites such as myo-inositol, fructose and sorbose also exhibited higher abundance at S2 in the unthinned control (Fig. 6). These compounds are known to accumulate in the vacuoles of cells within the fruit (Bae *et al*., 2014). The increased accumulation of these solutes in the cell’s vacuole could be an attempt to increase the osmotic concentration and influx of water supply, as water transport to fruit cells is critically important for early fruit development (Shiratake and Martinoia, 2007).

At S2, the unthinned control demonstrated lower sorbitol and higher fructose abundance than in fruit from the thinned treatment (Fig. 6). Sorbitol is the main sugar transported throughout the phloem in peach and is readily converted to fructose by SDH (Zhang *et al*., 2004; Morandi *et al*., 2008). Under heavy crop load conditions, fruit are in high competition for photosynthates (Grossman and DeJong, 1995). This limited carbon condition may explain why lower levels of sorbitol and higher levels of fructose are present in the fruit from the unthinned controls at S2 (Fig. 6), as fruit are perhaps rapidly converting sugar-alcohols into saccharides to be used for metabolic processes integral for survival under high carbon stress conditions.

Quinic acid is significantly higher in abundance in fruit from the carbon sufficient treatment at S2 and S3 (Fig. 6K), although levels drop before harvest (S4) in both the carbon sufficient and starved treatments, agreeing with previous research (Wu *et al*., 2005). Quinic acid is a product of the shikimic pathway, which initiates with imported carbohydrates (Walker and Famiani, 2018). The pentose phosphate pathway (PPP) converts available carbohydrates to glucose-6-phosphate (G6P), which are converted to erythose-4-phosphate (E4P). Phosphoenolpyruvate and E4P are then synthesized to form 3-deoxy-D-arabino-heptulosonic acid 7-phosphate (DAHP), which is then converted to dehydroquinic acid (DHQ). Quinate dehydrogenase subsequently converts DHQ to quinic acid, which can be used for the synthesis of chlorogenic acids (conjugates of quinic and cinnaminic acids; Guo *et al*., 2014). Chlorogenic acids are a group of phenolic acids (Clifford, 2000), which are important precursors for the development of anthocyanins and numerous other metabolites associated with quality attributes such as flavor, color, taste and nutrition (Walker and Famiani, 2018). This pathway is also responsible for the connection between the primary metabolism and the biosynthesis of various secondary metabolites, such as aromatic amino acids, folates, quinones, phytohormones, alkaloids, indole glucosinolates, flavonoids, hydroxycinnamic acids and lignins (Lara *et al*., 2020).

In fruits, the phenolic content is mostly comprised of phenolic acids and flavonoids (Häkkinen *et al*., 1999). Similar to the elevated levels of quinic acid (an intermediate for phenolic acids), levels of flavonoids such as catechin and tocopherol, were also higher in the carbon sufficient treatment (Figs. 6, 8). Thus, our data suggest that when carbon is in sufficient supply (achieved via thinning), fruit metabolism is able to prioritize the synthesis of secondary metabolites that help protect the fruit against abiotic/biotic stresses, pathogens and increase its’ quality attributes (Ullah *et al*., 2017). This prioritization of secondary metabolism under sufficient carbon supply has previously been shown in kiwifruit (Nardozza et al. 2019), where a sufficient carbon supply resulted in a higher abundance of anthocyanins in fruit flesh, when compared to a carbon starved (i.e. girdled) treatment. Few differences were observed in metabolite abundances between the thinned treatment and the unthinned control at harvest, when controlling for equal maturity. This may be due to the high level of priority given to the primary metabolism, even under stressed conditions (i.e. high fruit-to-fruit competition for photosynthates). When secondary metabolites and pathways are considered, the carbon sufficient treatment (thinned) shows elevated abundances (especially at S2), as a potential “luxury” investment, given the increased carbohydrate availability. Under these conditions, increased carbon can fuel the primary metabolism, which contributes to the synthesis of intermediates that act as precursors in the secondary metabolism to develop increased secondary metabolite content that results in enhance quality characteristics at harvest (Pott *et al*., 2019). Therefore, we hypothesize that metabolic differences observed as a response to carbon starvation/sufficiency may be more pronounced in the fruit’s secondary metabolism than in the central carbon (i.e. primary) metabolism at harvest when controlling for maturity, this should be further investigated in future studies.

### Catechin and a classified unknown metabolite may act as primers for superior or inferior fruit quality

When evaluating the metabolite profiles between the carbon sufficient and starved treatments at harvest (S4), only two compounds exhibited significant differences (Supplementary Table S4). These included a classified unknown, A244002 (Hummel *et al*., 2007), which was elevated under carbon starvation, and catechin, a flavonoid that was elevated under carbon sufficiency (Fig. 8A, D).

The classified unknown, A244002 has yet to be fully annotated, but it has been found previously in two legume plant metabolomes, alfalfa (*Medicago sativa*) and *Lotus japonicus* (Fester *et al*., 2011; 2014). In both occurrences, it appears that A244002 was associated with stress, elevated in plants forming mycorrhizal associations in alfalfa and increased in *Lotus japonicus* under depreciated soil nitrogen levels (Fester *et al*., 2011; 2014). There may be a potential link between this compound and a stress response in peach as well, as it remains significantly higher during carbon stress throughout development (Fig. 8D). Furthermore, when assessing how this compound relates to quality parameters at harvest, it demonstrates a significant negative correlation with both DMC and SSC (Fig. 8E, F).

Catechin is a phenolic compound that is found throughout several plant species, including many tree fruits like apricots, apples and cherries (Bernatoniene and Kopustinskiene, 2018) and has been reported as the most abundant flavan-3-ol in peach, especially in the peel (Chang *et al*., 2000). Phenolic compounds contribute to color, pigment synthesis, flavor and demonstrate several health benefits (Pietta *et al*., 1998). Catechin, like many antioxidants, helps to provide protection against plant pathogen pressure and abiotic stresses (Zhang *et al*., 2016). Higher levels of catechin were reported in peach exocarp when fruit was grown at higher altitudes, and were linked with superior quality (Karagiannis *et al*., 2016). Additionally, in a cultivar trial the highest levels of catechin were observed in peaches that also had the highest levels of SSC (Saidani *et al*., 2017). Therefore, our results suggest that under sufficient carbon supply, the tree is investing more resources for the synthesis of compounds for reproductive organs, which result in enhanced quality. This is supported by the significantly higher abundance of catechin observed in the carbon sufficient treatment (thinned) throughout development and at harvest (Fig. 8A). Furthermore, catechin has a strong positive correlation with both DMC and SSC at harvest, indicating enhanced quality (Fig. 8B, C).

Additionally, while significant differences in abundances for each of these compounds were observed across the developmental stages, the high abundance of both compounds at S2 could have a significant impact on fruit quality at harvest (i.e. DMC and SSC; Fig. 8). Therefore, these two compounds may act as respective primers early in development, providing a potential link between primary/secondary metabolism and fruit quality characteristics in peach.

## Conclusion

This study sought to determine the true impact of crop load, a critical preharvest factor, on peach fruit quality and metabolism. As quality and metabolism are heavily influenced by maturation, the use of Vis-NIRS technology allowed for sampling of equal maturity fruit. This novel approach enabled comparisons between differing carbon competition conditions (i.e. thinning severities), without the confounding influence of maturation. Physiological analyses showed that 2.6 fruit · cm^**-**2^ of TCSA could be a potential optimal crop load for ‘Cresthaven,’ to maximize quality without sacrificing excessive yields. Furthermore, when fruit of equal maturity coming from thinned trees (15 cm fruit spacing) were compared to the unthinned control, superior quality enhancements were noted, underscoring the true impact of crop load on fruit internal quality. Minimal differences in the fruit ionome were detected between treatments, although concentrations appeared to decrease with fruit development. Similarly, peach metabolite profiles appear to be heavily regulated by development and maturation. Several significant differences in metabolites were detected early on, with vast metabolic shifts in S2, when phenotypic differences are absent. Inversely, at S4, minimal differences in the fruit metabolome were observed, while vast differences in fruit quality and phenotype were prevalent. This contrasting trend between the metabolome and fruit quality indicate a potential metabolite priming effect for fruit quality development, as a result of variable carbon conditions. When photosynthates are not limited due to competition, fruit appear to begin prioritizing the synthesis of secondary metabolites related to internal fruit quality attributes at harvest. Only two metabolites were significantly different between treatments and strongly correlate to fruit quality (e.g. DMC and SSC) at harvest. Catechin and A244002 may act as potential primers in the peach fruit metabolome, providing a link between metabolite abundances and fruit quality enhancement or detriment, respectively.

## Supplementary data

Supplementary data are available at *JXB* online.

Table S1. Effect of thinning severity on peach leaf mineral composition.

Table S2. Temperature program from Titan MPS peach mesocarp microwave digestion.

Table S3. Element concentrations in peach mesocarp under carbon sufficient/starved conditions throughout development.

Table S4. Relative abundances of the 36 annotated metabolites in peach mesocarp under carbon sufficient/starved conditions throughout development.

Fig. S1. Fruit size distribution by thinning treatment.

Fig. S2. Principal component analysis biplot of variable carbon supply on peach fruit ionome.

Fig. S3. Heat map profiles of peach ionome across development between thinning treatments.

Fig. S4. Butanoic acid metabolite abundance across treatments throughout fruit development.

Fig. S5. Sorbose metabolite abundance across treatments throughout fruit development.

## Data availability statement

All data supporting the findings of this study are available within the paper and within its supplementary materials published online.

### Acknowledgments

We would like to especially thank Dr. Fernando Blanco-Cipollone, Mr. David Sterle, Ms. Emily Dowdy and Mr. Bryan Braddy for field data collection assistance and orchard management. Ms. Kat Chen is acknowledged for assistance with fruit photographs editing. The present article was supported by the Specialty Crop Block Grant Program of the U.S. Department of Agriculture (USDA) in cooperation with the Colorado Department of Agriculture (CDA) and the project #20202690 *‘Management strategies to maximize Colorado peach orchards productivity and fruit quality potential’*. Its contents are solely the responsibility of the authors and do not necessarily represent the official views of the USDA or CDA.

## Author Contributions

Conceptualization, I.S.M., B.M.A., J.M.C., J.E.P.; methodology, B.M.A., J.M.C., I.S.M.; software, B.M.A., J.M.C., I.S.M.; validation, B.M.A., I.S.M., J.M.C., J.E.P.; formal analysis, B.M.A., I.S.M.; investigation, B.M.A., I.S.M.; resources, I.S.M., J.E.P.; data curation, B.M.A., I.S.M.; writing-original draft preparation, B.M.A., I.S.M.; writing-review and editing, B.A., I.S.M., J.M.C., J.E.P.; visualization, B.M.A., I.S.M.; supervision, I.S.M.; project administration, I.S.M.; funding acquisition, I.S.M. All authors have read and agreed to the published version of the manuscript.

